# Subtle variations in a client protein determine bacterial Hsp90 dependence

**DOI:** 10.1101/2025.07.01.662558

**Authors:** Marie Corteggiani, Amine Ali Chaouche, Miha Bahun, Flora A. Honoré, Deborah Byrne, Sébastien Dementin, Mathieu E. Rebeaud, Olivier Genest

## Abstract

Chaperones ensure protein homeostasis and are conserved across species. The ATP-dependent chaperone Hsp90 is present from bacteria to eukaryotes, where it stabilizes and activates a wide range of substrate proteins called clients. However, what determines whether a protein depends on Hsp90 remains an open question. Here, we focused on the bacterial chaperone Hsp90 and its obligate client TilS (referred to as TilS_So_) in the bacterium *Shewanella oneidensis*. Although Hsp90 is indispensable in *S. oneidensis* under heat stress by protecting the essential protein TilS_So_ from degradation by the protease HslUV, Hsp90 is dispensable in *Escherichia coli*, suggesting that *E. coli* TilS (TilS_Ec_) is Hsp90 independent. We therefore compared the TilS orthologs with respect to *in vitro* stability, *in vivo* degradation, and interaction with Hsp90 to identify the determinants of the Hsp90 dependence. We found that in contrast to TilS_So_, TilS_Ec_ was more stable, was not degraded by protease in the absence of Hsp90, and did not interact with Hsp90, indicating that TilS_Ec_ is not a client of Hsp90. Chimeras between TilS_So_ and TilS_Ec_ as well as directed mutagenesis revealed a region of TilS_So_ that is key for protease degradation and Hsp90 protection. Consistent with these results, the growth of *S. oneidensis* producing TilS_Ec_ was no longer dependent on Hsp90 under heat stress. Conversely, Hsp90 became essential for the growth of *E. coli* that produced TilS_So_ instead of TilS_Ec_. Taken together, these results provide new insights into the mechanism of client protection by Hsp90 and the interplay between chaperones and proteases.

## Introduction

Chaperone proteins are present in all organisms to assist other proteins in folding, unfolding, preventing aggregation, disaggregation, and in some cases, degradation [1–3]. How chaperones identify and interact with these proteins (also called clients) remains an open question. Some chaperones interact with hydrophobic regions of the clients that are exposed during the folding process before being buried in the native conformation, thereby preventing protein aggregation, and promoting folding towards the native conformation. One of the most studied chaperones, Hsp70 (DnaK in prokaryotes) interacts with a stretch of hydrophobic residues exposed in virtually all unfolded proteins, although it can also target proteins later in the folding process [4–7]. For other chaperones such as Hsp90, no consensus sequence in clients is known. However, client features, such as hydrophobicity or intrinsically disordered regions, could direct Hsp90 recognition, and Hsp90 is believed to act late in the folding process [8–11].

Hsp90 is a highly conserved ATP-dependent chaperone in bacteria and eukaryotes [9,12–15]. It is a dimeric three-domain protein whose activity is regulated by large conformational changes induced by ATP binding and hydrolysis [13]. The N-terminal domain interacts with nucleotides, the middle domain mainly binds most of the clients, and the C-terminal domain allows dimerization. Several cochaperones, found only in eukaryotes, interact with Hsp90 to control and modulate its functional cycle depending on the client [2,16]. In both eukaryotes and in bacteria, Hsp90 functions in concert with the DnaK/Hsp70 chaperone system through direct interactions between Hsp70/DnaK and Hsp90 [12]. Although many clients of eukaryotic Hsp90 have been identified, the determinants that render a protein Hsp90-dependent are not fully understood and additional studies with other clients are needed.

*Shewanella oneidensis* is an aquatic gram-negative bacterium with a strong ability to adapt to environmental changes and stresses [17]. For example, it can support variations in a wide range of temperatures. Interestingly, we have shown that the Hsp90 chaperone is required in this bacterium to support growth under heat stress, making *S. oneidensis* a powerful model to study the Hsp90 chaperone [18]. Using a genetic selection, we have identified the client of Hsp90, TilS, which is responsible for Hsp90 essentiality at high temperature [18]. We found that Hsp90 protects TilS from degradation by HslUV, a conserved chaperone-protease machinery (**Figure 1A**) [19]. We demonstrated that direct binding between DnaK and Hsp90 is required to enable TilS protection and activation, suggesting a transfer of TilS from DnaK to Hsp90 and showing the interplay between chaperones and protease for post-translational TilS control [19,20].

**Figure 1:**
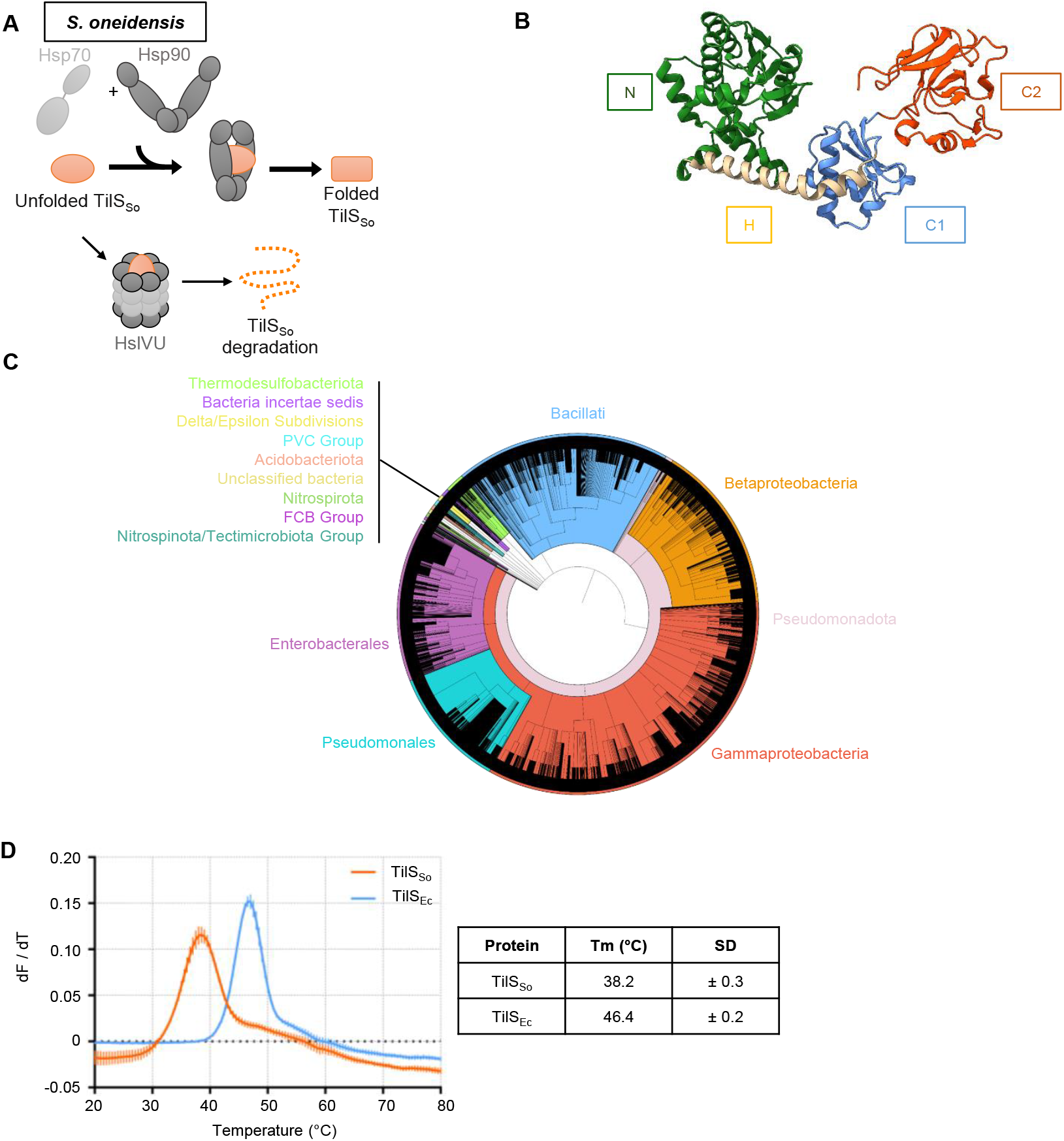
Generalities on the TilS protein. (A) Simplified hypothetical model of TilS_So_ requirement for Hsp90. In *S. oneidensis*, Hsp90 in collaboration with Hsp70/DnaK, is essential to promote TilS_So_ folding and thereby prevent its degradation. (B) Structure of TilS_Ec_ (PDB: 1NI5). Four domains are identified: the N-terminal (N) domain in green, the Helical (H) domain in gold, the C-terminal 1 (C1) in light blue, and the C-terminal 2 (C2) in orange. (C) Taxonomic tree of bacterial taxa based on TilS sequences. This circular taxonomic tree represents the taxonomic relationships between the different bacterial groups that possess TilS. Colors highlight the main bacterial clades, including Bacillati (blue), Betaproteobacteria (orange), Gammaproteobacteria (red), Pseudomonadales (cyan), Enterobacterales (purple), Pseudomonadota (light red) as well as other minor bacterial groups. (D) Determination of melting temperatures of TilS_So_ and TilS_Ec_ by Thermal Shift Assay. TilS_So_ or TilS_Ec_ were labeled with SYPRO Orange and incubated with increasing temperatures. The protein unfolding curves of three replicates from three independent experiments are shown as mean ± SD of derivatives of the fluorescence. The table indicates the melting temperatures as mean ± SD.

TilS is a conserved enzyme that modifies a tRNA to allow translation of the rare AUA codon into isoleucine, making it essential in bacteria [21–24]. TilS consists of an N-terminal catalytic domain, a long α-helix linker, and two C-terminal subdomains (**Figure 1B**). The mechanism of TilS activity has been elegantly demonstrated and involves large conformational changes: the two C-terminal subdomains hold and lock the tRNA while the N-terminal domain specifically catalyzes the addition of a lysine at cytidine 34 of the anticodon loop, resulting in a lysidine and switching the tRNA from methionine-specific to isoleucine-specific [24].

Surprisingly, although (i) TilS is essential in *Escherichia coli* [22], and (ii) Hsp90 is required for the protection and activation of TilS in *S. oneidensis* under heat stress [18], an *E. coli* strain that does not produce Hsp90_Ec_ is only slightly affected by the absence of Hsp90 under heat stress [25]. One hypothesis to explain these observations is that TilS from *E. coli* can reach its native conformation without the action of Hsp90, and therefore is not a client of Hsp90. If this hypothesis is correct, comparing the homologous TilS proteins from *E. coli* and *S. oneidensis* with respect to Hsp90 dependence could provide valuable insight into client recognition and specificity by Hsp90, as well as the interplay between Hsp90 and the HslUV protease.

Here, by comparing TilS orthologs from *E. coli, S. oneidensis* and other bacteria, we provide insight into the determinants that render TilS dependent or independent of Hsp90. Using *in vitro* approaches, chimera constructions, and *in vivo* phenotypes, we found that the degradation-sensitive TilS proteins, including TilS from *S. oneidensis* (TilS_So_), require Hsp90 for their stabilization. In contrast, *E. coli* TilS (TilSE_c_) did not require Hsp90 for protection, and therefore did not interact with Hsp90. In addition, we mapped a region of *S. oneidensis* TilS (TilS_So_) that is responsible for protein degradation by protease and protection by Hsp90. Taken together, these results provide a better understanding of the mechanism of client recognition by chaperones and proteases.

## Results

### Comparison of the TilS proteins from *E. coli* and *S. oneidensis*

A taxonomic tree based on 10500 TilS proteins distributed over more than 6000 different bacteria indicated that TilS is present in most bacteria and that its evolution follows bacterial phylogeny (**Figure 1C**). We focused on the TilS proteins from *E. coli* (TilS_Ec_) and from *S. oneidensis* (TilS_So_). The two TilS proteins share 35% sequence identity, TilS_So_ (464 amino acids, 52.1 kDa, Uniprot ID: Q8EGF9) is larger than TilS_Ec_ (432 amino acids, 48.2 kDa, Uniprot ID: P52097), and the key catalytic residues (including D130, E133 and R203, *S. oneidensis* numbering) are conserved [24,26] (**Figure S1A**). The N-terminal domain and the helical linker present a higher level of conservation (46 % identity) than the C1 and C2 domains (28 % identity). Additional amino acids are found at the N-terminal extremity (10 amino acids) and in the two C-terminal domains of TilS_So_ (**Figure S1A**). A comparison of the X-ray crystal structure of TilS_Ec_ (PDB 1NI5) with the AlphaFold model of TilS_So_ (AF-Q8EGF9) is shown in **Fig. S1B** [27].

To experimentally compare the two proteins, TilS_So_ and TilS_Ec_ were produced in *E. coli* and were purified. A small shift in size exclusion chromatography elution profiles was observed, in agreement with the low molecular mass difference of the two proteins (**Figure S1C**). Interestingly, thermal shift assays revealed that while the melting temperature (Tm) of TilS_Ec_ was 46.4°C, it was only 38.2°C for TilS_So_ (**Figure 1D**). This large difference in Tm values indicates a higher stability of TilS_Ec_ compared to TilS_So_, consistent with the hypothesis of a stronger requirement of chaperones for TilS_So_ compared to TilS_Ec_ under heat stress. In the same line of thought, we found that TilS_So_ was more susceptible to trypsin degradation than TilS_Ec_, again suggesting that TilS_So_ is less stable than TilS_Ec_ (**Figure S1D**).

### TilS_Ec_ is Hsp90-independent in contrast to TilS_So_

Since we have previously shown that TilS_So_ is a client of Hsp90 and that it interacts with Hsp90 [18], we wondered whether TilS_Ec_ behaves similarly as TilS_So_ or not. We first conducted bacterial two-hybrid experiments [28]. In these assays, the Hsp90 proteins from *E. coli* (Hsp90_Ec_) and from *S. oneidensis* (Hsp90_So_), as well as the TilS proteins from *E. coli* (TilS_Ec_) and from *S. oneidensis* (TilS_So_) were produced from plasmids as fusion proteins with the T18 or T25 catalytic domains of the adenylate cyclase from *Bordetella pertussis*. Combinations of plasmids were introduced in an *E. coli* strain deleted of the gene encoding the adenylate cyclase. If Hsp90 and TilS interact, the two catalytic domains are in proximity, reconstituting the enzymatic activity of the adenylate cyclase, leading to cAMP production and in turn to β-galactosidase activity. As already shown, we observed that TilS_So_ interacted with Hsp90_So_ (**Figure 2A**). TilS_So_ also interacted with Hsp90_Ec_, indicating that TilS_So_ is recognized by both Hsp90_Ec_ and Hsp90_So_. In contrast, background levels of β-galactosidase activity were measured when TilS_Ec_ was produced with Hsp90_Ec_ or Hsp90_So_, similarly to the levels measured when T18-TilS_Ec_ was produced with the T25 domain alone. These results indicate that Hsp90 does not interact with TilS_Ec_. As positive controls, we found that the Hsp90 fusion proteins retained their ability to dimerize since production of the T18 and T25 fusion proteins containing Hsp90_So_ or Hsp90_Ec_ led to a strong level of β -galactosidase activity. Finally, Western blot analysis indicated that the T18-TilS_Ec_ and T18-TilS_So_ fusion proteins were produced (**Figure S2A**).

**Figure 2:**
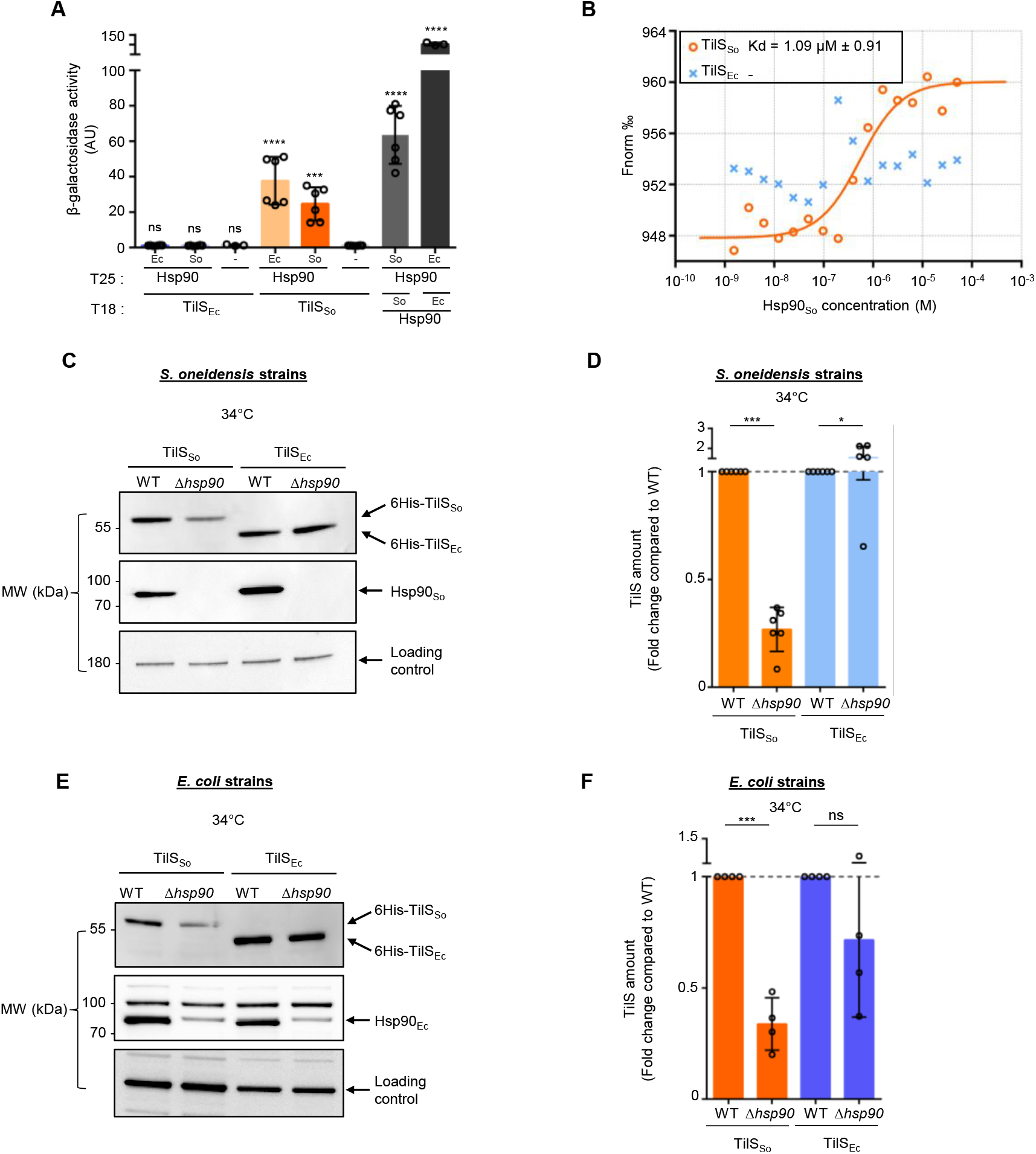
TilS_Ec_ is not a client of Hsp90. (A) Bacterial two-hybrid experiments showing the interaction between TilS and Hsp90. Hsp90_So_ and Hsp90_Ec_ were fused to the T25 domain of *B. pertussis* adenylate cyclase, and TilS_So_ and TilS_Ec_ were fused to the T18 domain. Bth101 *Δhsp90 E. coli* strains were transformed with two plasmids allowing the production of T18 and T25 fusions, respectively. The interaction between TilS and Hsp90 was monitored indirectly by following β-galactosidase activity. In negative control (−), the T25 domain alone was used, and in positive control the T18 domain was fused to Hsp90_Ec_ or Hsp90_So_ to monitor the interaction of Hsp90 dimer. Data from at least three replicates are shown as mean ± SD. (B) Microscale thermophoresis experiment to determine the interaction between Hsp90 and TilS_So_ or TilS_Ec_. 85 nM of TilS labeled with RED-tris-NTA 2nd generation dye (NanoTemper) was added to serial 2-times dilutions of Hsp90_So_ up to 50 µM. Fluorescence at 670 nm was measured, and values at 2.5 s were used to determine K_d_. The curves shown are representative of three independent experiments for TilS_So_ and two for TilS_Ec_. The mean of the Kd with standard deviation from three independent experiments is indicated. (C) Western blot showing the amount of TilS_So_ or TilS_Ec_ in *S. oneidensis*. The strains WT or *Δhsp90* with plasmids allowing the production of TilS_So_ or TilS_Ec_ with hexahistidine tag were grown at 34°C (heat stress conditions) and 0.02% arabinose was added. Two hours later, samples were analyzed by Western blot using anti-His antibody. Anti-Hsp90_So_ antibody was used as a control, and a contaminating band revealed by the anti-AtcJ antibody was used as a loading control. (D) Quantification of TilS from the Western blots shown in *C*. The peaks corresponding to the pixels of each band were quantified using ImageJ software. The amount of TilS_So_ or TilS_Ec_ in WT strain was set to 1, and each WT is compared to *Δhsp90* producing the corresponding TilS_So_ or TilS_Ec_ respectively. Data from six replicates are shown as mean ± SD. (E) Western blot showing the amount of TilS_So_ or TilS_Ec_ in *E. coli* in the same conditions as in *C*. Anti-Hsp90_Ec_ was used as control. The band above Hsp90_Ec_ is a contaminant detected by the anti-Hsp90_Ec_ antibody. In *C* and *E*, the Western blots shown are representative of at least four independent experiments. (F) Quantification as explained in *D* of the anti-His Western blot shown in *E*. Data from four replicates are shown as mean ± SD. In *A, D* and *F*, results of one-way ANOVA indicate whether the differences are significant (\*\*\**P* ≤ 0.001, \**P* ≤ 0.05) or not (ns, *P* > 0.05).

To confirm the results observed by two-hybrid and obtain binding parameters between the purified TilS proteins and Hsp90, we performed microscale thermophoresis (MST) experiments. This approach quantifies interactions by measuring the intensity variations of the fluorescently labeled proteins along a microscopic temperature gradient in the presence of increasing concentrations of an unlabeled partner protein. TilS_Ec_ and TilS_So_ were first labeled with RED-tris-NTA fluorescent dye that specifically interacts with the 6-His tag of the two proteins. In the MST experiments, the concentration of the RED-tris-NTA labeled TilS_so_ or TilS_Ec_ was kept constant (85 nM), while the concentration of the non-labeled binding partner Hsp90_So_ was varied between 0 µM-50 µM. For TilS_So_, the signal was fitted as a one-site binding curve indicating that the two proteins interact with a K_d_ of 1.09 µM (**Figure 2B and S2B**). As also shown by bacterial two-hybrid experiments, TilS_Ec_ did not interact with Hsp90 since no variation in fluorescence between the two proteins was observed by MST under these conditions (**Figure 2B and S2B**).

We have previously shown that the amount of TilS_So_ is dramatically reduced under heat stress in the absence of Hsp90 in *S. oneidensis* due to the activity of the HslUV protease [19]. To test whether TilS_Ec_ is also degraded in the absence of Hsp90, TilS_So_ or TilS_Ec_ with a 6-His tag at the N-terminal extremity was produced from an inducible promoter in the WT and Δ*hsp90 S. oneidensis* strains grown under heat stress (34°C), and the amount of the TilS protein was quantified by Western blot. Although the amount of TilS_So_ was reduced to about 30% in the Δ*hsp90* strain as expected, no reduction in the amount of TilS_Ec_ was observed in the absence of Hsp90 (**Figure 2C and D**). Interestingly, similar results were observed in *E. coli* WT or Δ*hsp90* strains (**Figure 2E and F**). These results show that in contrast to TilS_So_ at 34°C, TilS_Ec_ is not susceptible to degradation in the absence of Hsp90 both in *S. oneidensis* and in *E. coli*. When grown at 28°C, the amount of TilS_So_ and TilS_Ec_ did not significantly vary in *E. coli* and *S. oneidensis* WT or Δ*hsp90* strains (**Figure S2C-F**).

Altogether, these experiments show that in contrast to TilS_So_, TilS_Ec_ does not interact with Hsp90 in our experimental conditions, and its amount is not reduced in the absence of Hsp90, strongly suggesting that TilS_Ec_ is not an Hsp90 client, whereas TilS_So_ is.

### TilS identity determines the Hsp90-dependent growth of bacteria

As proposed above, TilS_Ec_ does not require the activity of the Hsp90 chaperone to be functional, unlike TilS_So_. It would imply that the growth defect of the *S. oneidensis* Δ*hsp90* strain under heat stress would be rescued by the production of TilS_Ec_ instead of TilS_So_. To test this hypothesis, *tilS*_*So*_ was replaced by *tilS*_*Ec*_ on the chromosome of the *S. oneidensis* MR1 WT and Δ*hsp90* strains using homologous recombination, leading to the strains Δ*tilS*_*So*_ *tilS*_*Ec+*_ and Δ*hsp90*_So_ Δ*tilS*_*So*_ *tilS*_*Ec*_^*+*^, respectively. We measured the growth of these strains as well as the WT and Δ*hsp90*_So_ strains that were incubated at either 28°C (the non-stress temperature at which Hsp90_So_ is dispensable) or 35°C (the heat stress temperature at which Hsp90_So_ is required). We first observed that TilS_Ec_ can functionally substitute for TilS_So_ to support growth of *S. oneidensis* since the four strains grew similarly at 28°C (**Figure 3A and S3A**), and under heat stress (35°C) the strain producing TilS_Ec_ instead of TilS_So_ in the presence of Hsp90 (Δ*tilS*_*So*_ *tilS*_*Ec*_^*+*^) grew as the WT strain (**Figure 3B**). As expected, the absence of Hsp90 led to a strong growth defect at 35°C (Δ*hsp90*_So_ vs WT), but not at 28°C (**Figure 3A and B**) [18]. Interestingly, replacing *tilS*_*So*_ by *tilS*_*Ec*_ in the *S. oneidensis hsp90*-knockout strain (Δ*hsp90*_So_ Δ*tilS*_*So*_ *tilS*_*Ec*_^*+*^) dramatically improved growth of the Δ*hsp90*_So_ strain at 35°C (**Figure 3B**). Similarly, the growth defect under heat stress of the Δ*hsp90*_So_ strain was rescued by the production of TilS_Ec_ instead of TilS_So_ when bacterial growth was performed on LB-agar plates (**Figure S3B and C**). Additional growth temperatures (34°C and 36°C) were also tested. Although at 34°C, the presence of TilS_Ec_ almost totally rescued the growth defect of the Δ*hsp90*_So_ strain, it only slightly rescued growth at 36°C (**Figure S3C and D**). This could suggest that additional proteins, besides TilS, that are important for bacterial growth are impacted by the absence of Hsp90 at temperatures above 35°C. In agreement with this hypothesis, the slight growth defect of the Δ*hsp90*_So_Δ*tilS*_*So*_ *tilS*_*Ec+*_ at 35°C was fully rescued by the production of Hsp90 from a plasmid (**Figure S3E**).

**Figure 3:**
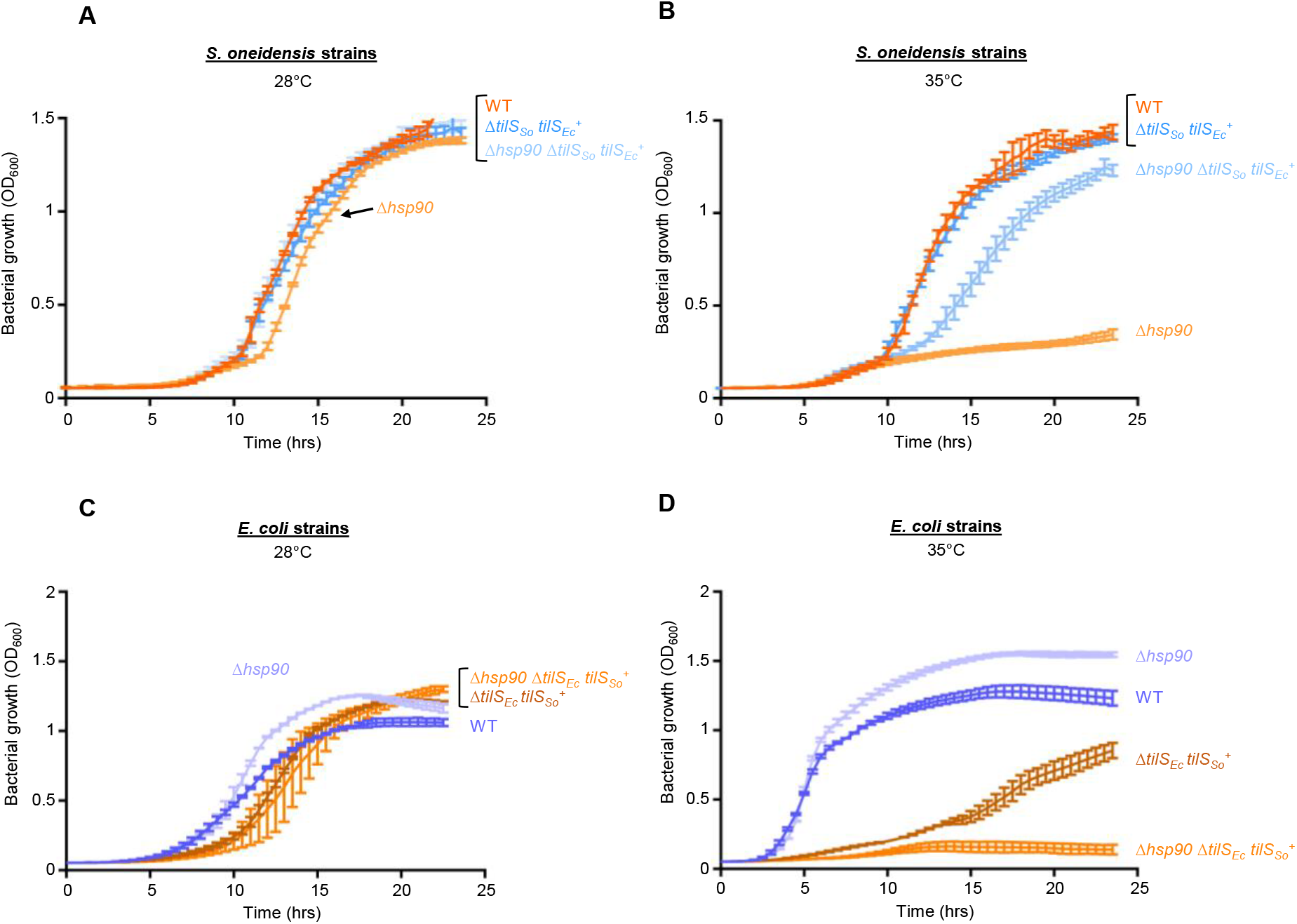
The identity of TilS dictates the dependence on Hsp90 in *S. oneidensis* and in *E. coli*. (A) Bacterial growth of *S. oneidensis* strains carrying *tilS* from *E. coli*. The strains *S. oneidensis* MR-1 wild-type (WT), deleted of *hsp90* (*Δhsp90*) with natural TilS or replaced by TilS from *E. coli* (Δ*tilS*_*So*_ *tilS*_*Ec*_^*+*^) were grown in microplates in LB medium at 28°C with shaking for 24 hours. (B) Bacterial growth of the strains used in *A* at 35°C. (C) Bacterial growth of *E. coli* strains carrying *tilS* from *S. oneidensis*. The strains *E. coli* MG1655 wild-type (WT), deleted of *hsp90* (*Δhsp90*) with natural TilS or replaced by TilS from *S. oneidensis* (Δ*tilS*_*Ec*_ *tilS*_*So*_ ^*+*^) were grown in microplates in LB medium at 28°C with shaking for 24 hours. (D) Bacterial growth of the strains used in *C* at 35°C. In *A, B, C* and *D*, data from three replicates are shown as mean ± SD.

In contrast to *S. oneidensis*, the absence of *hsp90* in *E. coli* does not lead to significant growth defects even under heat stress [25]. Since we know that Hsp90 is essential for TilS_So_ stabilization at temperatures above 35°C, we wondered whether we could transform *E. coli* from an Hsp90-independent strain to an Hsp90-dependent strain only by exchanging its TilS (*i*.*e*. TilS_Ec_) for TilS_So_. The *tilS* gene in *E. coli* MG1655 WT and Δ*hsp90*_Ec_ was replaced by *tilS*_*So*_ by homologous recombination leading to strains Δ*tilS*_*Ec*_ *tilS*_*So*_^*+*^ and Δ*hsp90*_*Ec*_ Δ*tilS*_*Ec*_ *tilS*_*So*_^*+*^, respectively. At 28°C, the four strains grew similarly (**Figure 3C**). However, at 35°C, the growth of Δ*tilS*_*Ec*_ *tilS*_*So*_^*+*^ was delayed compared to WT, indicating that TilS_So_ cannot fully compensate for the absence of TilS_Ec_. Nevertheless, we found that deleting *hsp90* dramatically reduced the growth of the *E. coli* strain that produces TilS_So_ instead of TilS_Ec_ (**Figure 3D**). Therefore, by exchanging the TilS proteins, *E. coli* was successfully converted to a strain whose growth depends on the Hsp90 chaperone. Similar results were observed when growth was performed on LB-agar plates (**Figure S3F and G**).

Altogether, these experiments demonstrate that the origin of the TilS protein (i.e. from *S. oneidensis* or from *E. coli*) dictates the dependence of a strain on the Hsp90 chaperone. Indeed, the growth defect of a *S. oneidensis* strain devoid of Hsp90 can be rescued by TilS_Ec_, a protein that does not need Hsp90 to be functional. Conversely, an *E. coli* strain that produces TilS_So_ requires Hsp90 for growth at 35°C, a temperature at which the TilS_So_ protein needs Hsp90 to be protected.

### The C-terminal domain of TilS_So_ is responsible for the dependence on Hsp90

We next wanted to identify the region of TilS_So_ that confers dependence on Hsp90. To this end, we constructed chimeras by swapping domains between TilS_So_ and TilS_Ec_ (**Figure 4A**). As described above, TilS is constituted by an N-terminal domain (N), a long alpha helix (H), and a C-terminal domain divided in the two subdomains C1 and C2 (**Figures 1B, S1A-B**). The chimeras were named according to the origin of each of their four domains, with “S” for *S. oneidensis* and “E” for *E. coli*. For example, “SEEE” stands for the chimera with the N-terminal domain of TilS_So_ and the three other domains of TilS_Ec_. The chimeras were produced from a pBad inducible promoter in the *S. oneidensis* WT and Δ*hsp90* strains grown at 28°C and 34°C. To quantify the amount of the TilS chimeras in the two strains, total proteins were separated on SDS-PAGE and the chimeras were detected by Western blot (**Figures 4B and C, and S4A and B**).

**Figure 4:**
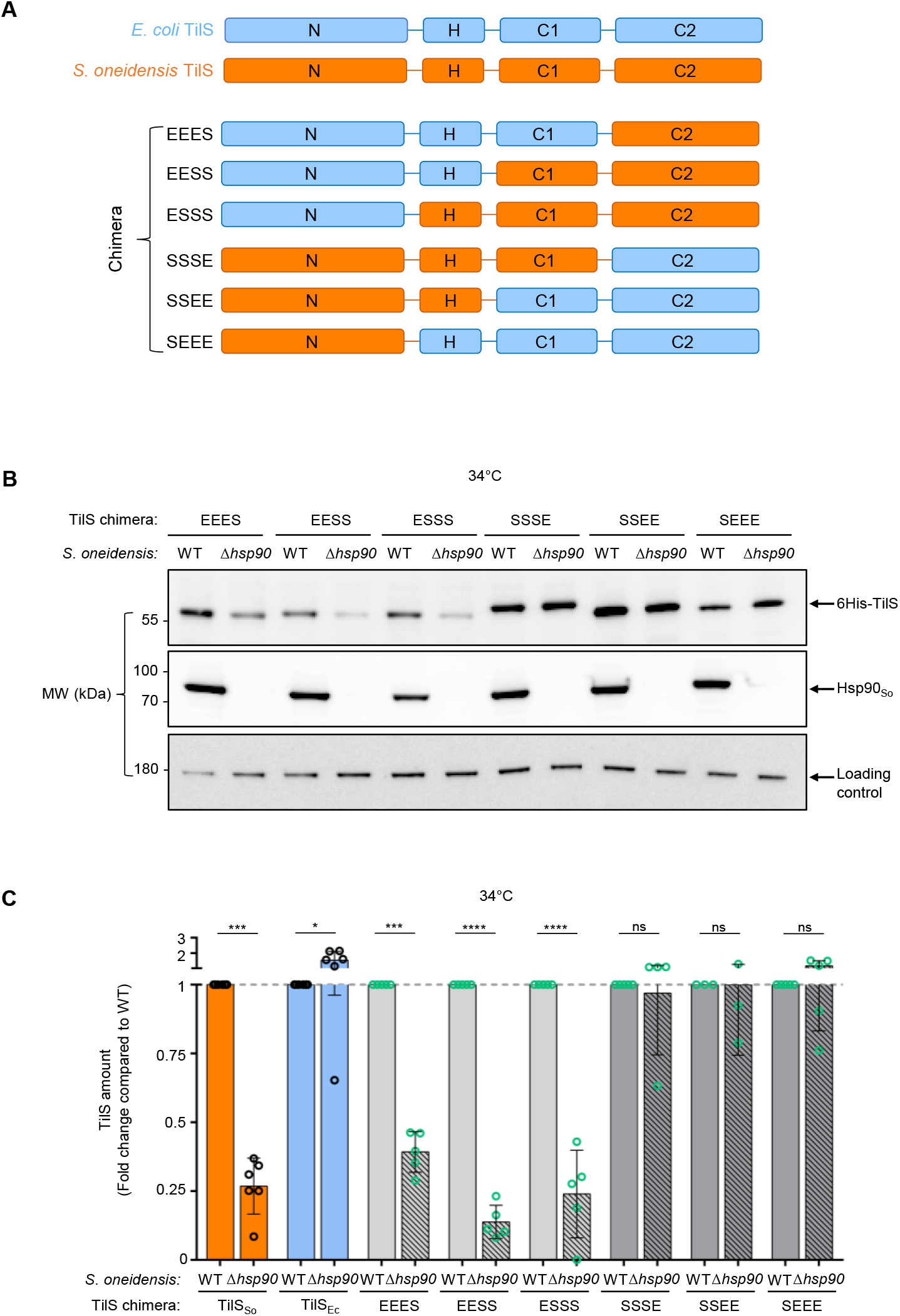
The C-terminal domain of TilS dictates Hsp90 dependence. (A) Schematic showing the organization in domains of TilS and how they were mixed to give 6 TilS_Ec_ – TilS_So_ chimeras. « E » corresponds to TilS domain from *E. coli*, and « S » to domain from *S. oneidensis*. (B) Western blot showing the abundance of TilS chimeras in *S. oneidensis* WT or Δ*hsp90*. The strains WT or Δ*hsp90* with plasmid pBad33 allowing the production of 6His-TilS chimeras were grown at 34°C, and 0.02% arabinose was added. After 2 hours, samples were analyzed by Western blot with anti-His antibody. Anti-Hsp90 was used as a control and Anti-AtcJ as neutral loading control. This Western blot is representative of at least three independent experiments. (C) Quantification of the Western blot shown in *B*. The peaks corresponding to the pixels of each band were quantified using ImageJ software. The amount of each chimera in WT strain was set to 1, and each WT is compared to *Δhsp90* producing the corresponding chimera respectively. Data from at least three replicates are shown as mean ± SD. Results of one-way ANOVA indicate whether the differences are significant (\*\*\*\**P* ≤ 0.0001, \*\*\**P* ≤ 0.001) or not (ns, *P* > 0.05).

Although, as indicated above, the amount of TilS_So_ is considerably reduced in the absence of Hsp90 (**Figures 2C, 2D, and 4C**), we found that replacing the single C2 domain of TilS_So_ with that of TilS_Ec_ (“SSSE” chimera) was sufficient to stabilize TilS_So_ in the absence of Hsp90. Exchanging the C1 and C2 domains (chimera “SSEE”) or H, C1, and C2 domains (chimera “SEEE”) of TilS_So_ by the domains of TilS_Ec_ also led to stabilization of the chimeras in the absence of Hsp90 (**Figure 4B and C**). Contrarily, although the amount of TilS_Ec_ was not reduced in the absence of Hsp90 compared to WT (**Figures 2C, 2D, and 4C**), substituting the C2 domain of TilS_Ec_ by the one of TilS_So_ (chimera “EEES”) was sufficient to lead to a lower level of chimera in the absence of Hsp90, therefore indicating a dependence on the Hsp90 chaperone (**Figure 4B and C**). Similar results, i.e., a lower amount of chimeras in the absence of Hsp90, were obtained with the chimeras “EESS” and “ESSS”.

These results strongly suggest that the C2 domain of TilS_So_ is responsible for the dependence on the Hsp90 chaperone. When Hsp90 does not protect it, this domain most likely directs TilS_So_ to degradation. Indeed, we found that Hsp90, which does not interact with TilS_Ec_ (**Figure 2A and B**) protects the “EEES” chimera, possessing only the C2 domain of TilS_So_, from degradation (**Figure 4**). Interestingly, the AlphaFold3 prediction of the binding between Hsp90_So_ and TilS_So_ positions the two proteins in interaction via the C2 domain of TilS_So_, while no binding was predicted with TilS_Ec_ (**Figure S5**).

### Identification of point mutations in the C2 domain of TilS_So_ that modify Hsp90 dependence

Based on our observations that TilS_So_ is an obligate client of Hsp90, while TilS_Ec_ is not, we aimed to predict Hsp90 essentiality in a given strain by analyzing its TilS protein. To this end, we hypothesized that TilS proteins evolutionary close to TilS_Ec_ would not require Hsp90 for stability, while those that are close to TilS_So_ would. To test this, a phylogenetic tree was built based on TilS C2 domains (**Figure S6**), and several TilS orthologs were selected and tested for their stability *in vivo* in *S. oneidensis*. However, we could not validate this hypothesis because some TilS proteins close to TilS_Ec_, such as TilS from Salmonella Typhymurium, were dependent on Hsp90 as shown with our *in vivo* degradation assay (**SI results, and Figure S7**). Interestingly, we were surprised to observe that TilS from *Shewanella xiamenensis* (TilS_Sx_), a protein with over 90% identity to TilS_So_, did not behave like TilS_So_ with regard to Hsp90 dependence. Indeed, we found that the amount of TilS_Sx_ was not reduced in the absence of Hsp90_So_, as observed with TilS_So_ (**Figures 5A, S7A, and S7C**). Comparison of the primary sequences between the two proteins revealed a few point mutations in the C2 domain, and in particular the L340 residue in TilS_So_ that is replaced by a proline residue in TilS_Sx_ (**Figure 5B**). We constructed the single mutant Hsp90_So_L340P to test whether this proline residue could be responsible for the stabilization of TilS_Sx_ in the absence of Hsp90. Interestingly, we found that, in contrast to the wild-type protein, the amount of the mutant protein TilS_So_L340P was only slightly reduced in the absence of Hsp90 (**Figures 5A and S8A**). Replacing L340 of TilS_So_ by residues other than proline (serine or glycine) (**Figure S8B**), or replacing the last 7 residues of TilS_So_ by the 8 last ones of TilS_Sx_ (**Figure S8C**) did not lead to stabilization of TilS_So_ in the absence of Hsp90. These results indicate that mutating the leucine 340 of TilS_So_ into a proline is crucial to stabilize the protein in the absence of Hsp90, as naturally found in TilS_Sx_.

**Figure 5:**
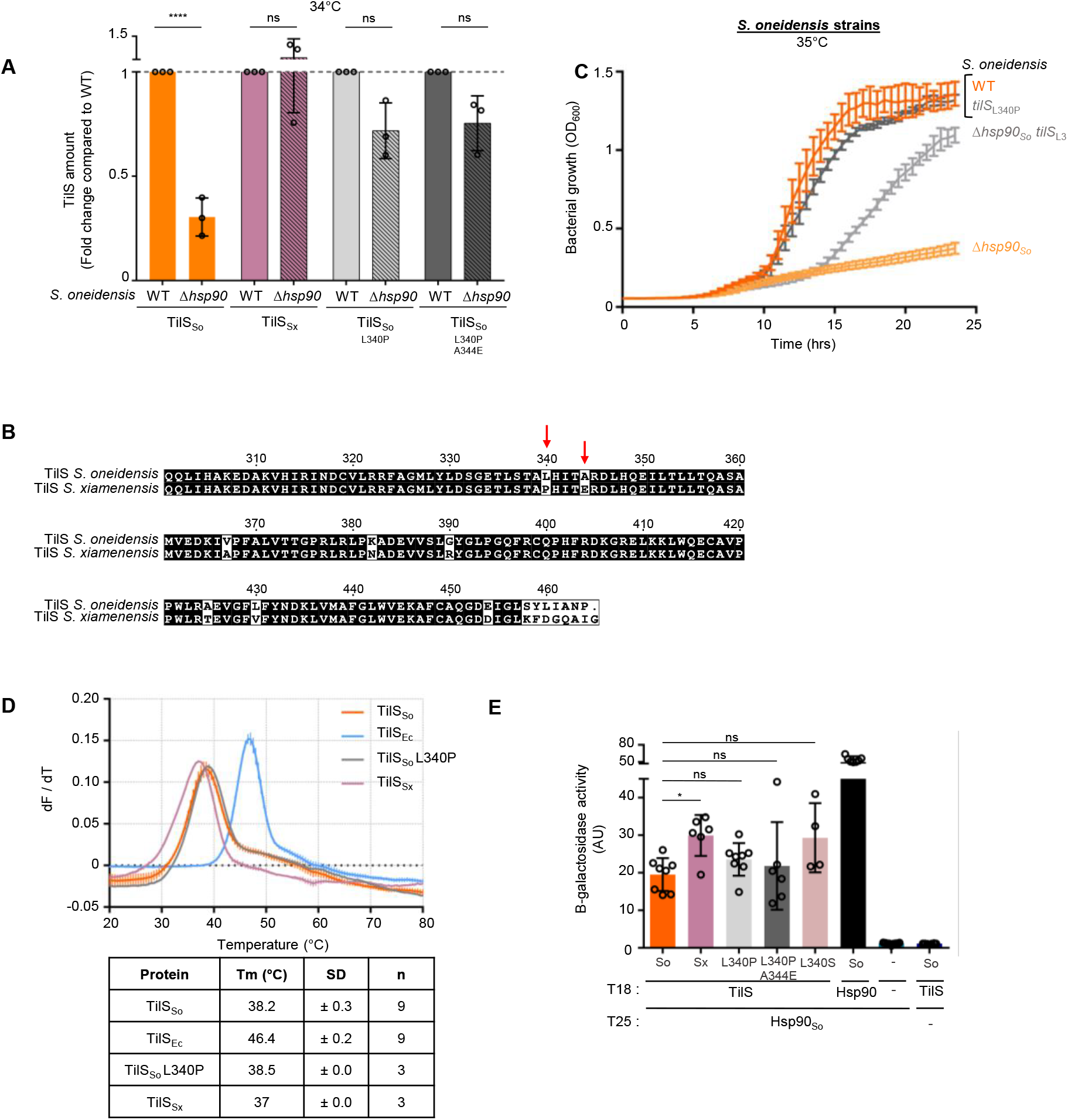
A single point mutation in TilS_So_ decreases Hsp90 dependence. (A) Graph showing the amount of TilS_Sx_, and TilS_So_ WT or mutants from the Western blot in Fig. S8A. The *S. oneidensis* WT or *Δhsp90* strains with plasmid allowing the production of TilS_So_ or TilS_Sx_ with hexahistidine tag were grown at 34°C (heat stress conditions) and 0.02% arabinose was added. Two hours later, samples were analyzed by Western blot using anti-His antibody. This graph shows the quantification of three independent Western blots. The peaks corresponding to the pixels of each band were quantified using ImageJ software. The amount of TilS_So_ or TilS_Sx_ in WT strain was set to 1, and each WT is compared to *Δhsp90* producing the corresponding TilS_So_ or TilS_Sx_ respectively. Data from three replicates are shown as mean ± SD. The results of one-way ANOVA indicate whether the differences are significant (\*\*\*\**P* ≤ 0.0001) or not (ns, *P* > 0.05). (B) Partial alignment of the sequences of TilS_So_ and TilS from *S. xiamenensis* (TilS_Sx_) using the multiple sequence alignment tool «Multalin» (http://multalin.toulouse.inra.fr/multalin/). TheESPript program (https://espript.ibcp.fr/ESPript/ESPript/) was used to display the alignment. Numbering of TilS_So_ and TilS_Sx_. The red arrows indicate the residues that were mutated. (C) Bacterial growth in liquid media of the *S. oneidensis* strains MR-1 wild-type (WT) or deleted of *hsp90* (*Δhsp90*) with the *tilS* gene WT or coding for the mutation L340P. The cells were grown in microplates in LB medium at 35°C with shaking for 24 hours. Data from six replicates are shown as mean ± SD. (D) Determination of the melting temperatures of TilS_So_ WT or L340P, TilS_Ec_, or TilS_Sx_ by Thermal Shift Assay. The TilS proteins were labeled with SYPRO Orange and incubated with increasing temperatures. The protein unfolding curves of at least three replicates are shown as mean ± SD of derivatives of the fluorescence. The table indicates the melting temperatures as mean ± SD. (E) Bacterial two-hybrid experiments showing interaction of TilS_So_ WT or mutants, or TilS_Sx_ with Hsp90_So_. Hsp90_So_ was fused to the T25 domain of *B. pertussis* adenylate cyclase and the five TilS proteins were fused to the T18 domain. Bth101 *Δhsp90* strains were transformed with two plasmids allowing the production of T18 and T25 fusions, respectively. The interaction between TilS and Hsp90 was monitored indirectly by following β-galactosidase activity. In the negative control (−), T25 or T18 domains alone were used, and in the positive control T25 and T18 domain were both fused to Hsp90_So_ to monitor the interaction of Hsp90 dimer. Data from at least three replicates are shown as mean ± SD. The results of one-way ANOVA indicate whether the differences are significant (\**P* ≤ 0.05) or not (ns, *P* > 0.05).

We then wondered whether the stabilization of TilS_So_ by the mutation L340P would allow bacteria to grow with a lower dependence on Hsp90 under heat stress. The corresponding mutation in *tilS*_*So*_ was introduced in the chromosome of *S. oneidensis* WT and Δ*hsp90*. In the WT background, the strain producing TilS_So_L340P instead of TilS_So_ grew similarly to the WT strain at 28°C and 35°C (**Figures 5C and S8D-F**). More importantly, production of TilS_So_L340P did partially rescue the growth defect of the *S. oneidensis* Δ*hsp90* strain (**Figure 5C**), confirming that TilS_So_L340P can be functional without the assistance of Hsp90.

We further characterized TilS_Sx_ and TilS_So_L340P to investigate why they can work independently of Hsp90, as was observed with TilS_Ec_. The proteins were purified and size exclusion chromatography elution profiles of the two proteins were similar to TilS_So_ (**Figure S9A**). Surprisingly, we found that TilS_Sx_ and TilS_So_L340P behaved more like TilS_So_ than TilS_Ec_ with regard to intrinsic protein stability and interaction with Hsp90. Indeed, thermal shift assays indicated that the Tm of TilS_Sx_ (37.0°C) and TilS_So_L340P (38.5°C) were closer to TilS_So_ (38.2°C) than TilS_Ec_ (46.4°C) (**Figure 5D**). Similarly, the trypsin digestion profiles of TilS_Sx_ and TilS_So_L340P resembled those obtained with TilS_So_ (**Figure S9B**). Finally, two-hybrid and MST experiments revealed that TilS_Sx_ and TilS_So_L340P interacted with Hsp90, as observed with TilS_So_, but in contrast to TilS_Ec_ (**Figure 5E and S9C**).

Altogether, based on the TilS ortholog from *S. xiamenensis*, we identified a single mutation, L340P, in TilS_So_ that switches the TilS protein from Hsp90-dependent (TilS_So_ WT) to Hsp90-independent (TilS_So_L340P). Since TilS_So_L340P still interacts with Hsp90, this suggests that the mutation prevents the degradation by the HslUV protease.

## Discussion

In this study, we aimed to explore the determinants that govern whether a protein is dependent on the bacterial Hsp90 chaperone. Our client model was the TilS protein, an essential enzyme whose function is to modify a tRNA. We have previously found that TilS is an obligate Hsp90 client under heat stress in the bacterium *S. oneidensis* [18]. Indeed, we have shown that Hsp90 protects TilS_So_ from degradation by the HslUV protease [18,19]. Since TilS is also essential in *E. coli*, but Hsp90 is dispensable for growth in this bacterium, we hypothesized that TilS_So_ and TilS_Ec_ do not have the same requirement for Hsp90. Therefore, we compared these TilS proteins. We found that although the two TilS orthologs share 35 % sequence identity and a common global fold (**Figure S1 A-B**), they do not behave similarly with respect to Hsp90 dependence. Indeed, in contrast to TilS_So_, whose level was dramatically reduced under heat stress in the absence of Hsp90, the level of TilS_Ec_ did not differ significantly between WT and Δ*hsp90* strains (**Fig 2C-F**). In addition, TilS_Ec_ did not interact with Hsp90, whereas TilS_So_ did (**Figure 2A-B**), suggesting that TilS_Ec_, unlike TilS_So_, is not a client of Hsp90. To more precisely define the regions of TilS that dictate Hsp90 dependence, chimeras between the two TilS proteins were constructed, allowing the identification of the C2 domain of TilS_So_ that is responsible for TilS degradation and Hsp90 dependence (**Figure 4**). Indeed, the amount of the chimera consisting of TilS_Ec_ with only the C2 domain of TilS_So_ (named “EEES”) was strongly reduced in the Δ*hsp90* strain, whereas TilS_So_ was stabilized by exchanging its C2 domain with that of TilS_Ec_ (named “SSSE”). In addition, we found that replacement of residue L340 of TilS_So_ with a proline was sufficient to partially abolish its degradation in the absence of Hsp90 (**Figure 5**). The mutation of this leucine residue to serine or glycine did not stabilize TilS_So_. This suggests that the local reorientation of the region near L340, or the increased rigidity of the protein induced by proline, is sufficient to prevent TilS_So_ degradation, rather than the nature of the leucine residue per se. Taken together, these experiments indicate that the C2 domain of TilS_So_ is key to triggering TilS_So_ degradation in the absence of Hsp90, and to allowing TilS to be protected by Hsp90.

Based on our results, we can propose a model in which Hsp90 interacts with the C2 domain of TilS_So_, thereby limiting the accessibility of the HslUV protease to an as yet unidentified degron probably located in the C2 domain of TilS_So_. After Hsp90-assisted folding of TilS_So_, the region recognized by HslUV could be buried and no longer accessible to the protease. This model is also supported by AlphaFold 3 predictions (**Figure S5**), which propose an interaction between a dimer of Hsp90_So_ and the C2 domain of TilS_So_. In these predictions, binding to Hsp90 occurs through the cleft located between the two monomers of Hsp90_So_, which is known to be important for binding to multiple clients [29]. Interestingly, mutations in this region of Hsp90 (W476R and L563A, Hsp90_So_ numbering) have been shown to prevent binding to clients, including TilS_So_ [18,29]. Alternative models obtained with AlphaFold 3 always position the C2 domain of TilS_So_ in contact with Hsp90_So_, but the other domains of TilS_So_ adopt different orientations without interacting with Hsp90 (**Figure S5B**). How Hsp90 interacts with its clients has long been of interest [8–11]. Studies performed with eukaryotic Hsp90 clients, including for example the Tau protein [30] and α-synuclein [31], suggest that hydrophobic patches lead to Hsp90-client stabilization. Recently, the laboratory of Brian Freeman identified over 1000 proteins that associated with Hsp90 in yeast using a cross-linking approach followed by mass spectrometry [11]. Mapping the chaperone binding sites in the clients revealed that Hsp90 preferentially interacts with intrinsically disordered regions of proteins. We used predictor tools including AIUPred algorithm to search for putative intrinsically disordered regions in the TilS_So_ sequence; however, no such regions were identified (**Figure S10A**) [32]. Interestingly, AlphaFold 3 indicates the presence of an unstructured loop in the C2 domain of TilS_So_, in the vicinity of residue L340, but this loop is not shown in the X-ray crystal structure of TilS_Ec_ (PDB 1NI5) (**Figure S10B**). While this may seem appealing, it must be treated with high caution since the structures of TilS_So_ and TilS_Sx_ shown here are only predictions and this loop could appear as an artefact.

The conclusions regarding the stability of TilS and its dependence on Hsp90 were supported by *in vivo* phenotypes. *E. coli* could be transformed from an Hsp90-dispensable strain to an Hsp90-essential strain by producing TilS_So_ instead of TilS_Ec_ (**Figure 3D**) [25]. Conversely, although *S. oneidensis* requires Hsp90 for growth under heat stress, we generated a *S. oneidensis* strain whose growth became Hsp90-independent only by replacing TilS_So_ with TilS_Ec_ (**Figure 3B**), or by introducing the single mutation L340P in TilS_So_ (**Figure 5C**). However, the restoration of the growth phenotype was almost lost when the temperature increased by only 1°C, although we know that TilS_Ec_ is functional at this temperature in *E. coli* (regular *E. coli* growth temperature is 37°C) (**Figure S3D**). Therefore, we can hypothesize that, although TilS_So_ is the only essential protein whose level dramatically decreases in the absence of Hsp90 at 35°C, the absence of Hsp90 at 36°C would impact several other proteins. In that case, compensating for the loss of TilS_So_ by producing TilS_Ec_ would not be sufficient to support growth of *S. oneidensis* because several other proteins or another essential protein would be inactivated by the increased temperature in the absence of Hsp90 (36°C).

The interplay between the Hsp90 chaperone and the HslUV protease has previously been documented [19,33,34]. Fauvet *et al*. proposed that aggregation-prone proteins are degraded by HslUV in an Hsp90-dependent process [34]. We have shown that TilS_So_ is degraded by the protease HslUV when it is not protected by Hsp90, and the severe growth defect of the *S. oneidensis* Δ*hsp90* strain was suppressed by deletion of *hslUV* [19]. Similarly, defects in colibactin toxin production in *E. coli hsp90* knockout strain were rescued by deletion of the gene encoding the HslV protease [33]. These observations suggest that the main function of Hsp90 is to protect its clients from protease degradation, most likely by facilitating their folding. The present study highlights differences in TilS orthologs regarding chaperone dependence and protease degradation. Based on protein intrinsic stability, we found that TilS_Ec_ and the TilS proteins from the *Shewanella* genus (TilS_So_ and TilS_Sx_) fell into two different categories. Indeed, thermal shift assays on purified proteins indicate that TilS_So_ and TilS_Sx_ are less stable than TilS_Ec_, with a shift higher than 8°C in the melting temperatures of the proteins from the two categories (**Figure 1D and 5D**). This result correlates with the fact that Hsp90 interacts with TilS_So_ and TilS_Sx_, but not with TilS_Ec_ (**Figure 2A-B, 5E, and S9C**). These observations are in agreement with a global study in yeast demonstrating that the intrinsically unstable state of kinases leads to recognition by Hsp90 [35]. An example is provided by the oncogenic v-Src kinase, an Hsp90-dependent protein, and the Hsp90-independent c-Src kinase, that share 98 % sequence identity [36]. c-Src was more stable than v-Src in thermal unfolding measurements, with a 4°C difference in the melting temperature of the two proteins, and v-Src was more prone to aggregation than c-Src.

Interestingly, additional differences were found in the *in vivo* stability of the TilS orthologs belonging to the *Shewanella* genus. Indeed, TilS_So_ was degraded by HslUV in the absence of Hsp90 [19], whereas TilS_Sx_ was not degraded (**Figure 5A**). Thus, despite conserving their ability to interact with Hsp90, these two proteins have evolved to be either protease sensitive (TilS_So_) or resistant (TilS_Sx_). This difference may be due to a single mutation, since mutating L340 to P in TilS_So_ is sufficient to stabilize it in the absence of Hsp90. Therefore, TilS_So_L340P and TilS_Sx_ are interesting proteins for studying the interplay between chaperones and proteases, as their behavior uncouples Hsp90 protective activity from HslUV proteolytic activity. The physiological relevance of TilS_Sx_ still interacting with Hsp90, despite not being degraded, remains unclear. It is possible that under certain stress conditions, this TilS protein requires the chaperone activity of Hsp90 to be functional.

These observations raise the question of the evolution of TilS in relation to its dependence on Hsp90. A taxonomic tree was constructed based on full-length TilS (**Figure 1C**) to assess the distribution of TilS, and a phylogenetic tree of the C2 domain of TilS (**Figure S6**). Although the evolution of TilS follows the bacterial phylogeny, we could not correlate the evolution of the C2 domain of TilS with a dependence on Hsp90 (**SI Results and Figure S7**). Indeed, as explained above, although the TilS proteins from the bacteria *S. oneidensis* and *S. xiamenensis* have more than 90 % sequence identity, TilS_So_ required Hsp90 for protection against protease degradation, while TilS_Sx_ did not. Interestingly, mutations may arise in the essential TilS protein because of the protection provided by the Hsp90 chaperone. Consistent with this, Hsp90 has been shown to play an important role in evolution in both eukaryotes and bacteria, and Hsp90 belongs to the family of “global modifiers” [37–41]. Hsp90 buffers the phenotypic effects of mutations that occur in proteins. Indeed, by stabilizing and allowing the folding of a mutated client, Hsp90 prevents the phenotype associated with that mutation from being expressed. These phenotypes can be revealed under conditions in which Hsp90 is titrated, for instance, by other clients.

The physiological relevance of the dependence on Hsp90 and/or degradation by protease regarding the cellular availability of the essential TilS protein is unclear, though it may play a significant role in regulating bacterial fitness in response to environmental stressors. For example, we can hypothesize that a less stable TilS may confer a fitness advantage to certain bacterial species. With the assistance of Hsp90, TilS_So_ may be better adapted than TilS_Ec_ to the likely more diverse environmental conditions of the ecological niche inhabited by *S. oneidensis*, compared to the relatively stable environment of the intestinal tract where *E. coli* resides [17].

In conclusion, our study on the TilS_So_ protein and some of its close orthologs highlights the variety of mechanisms implemented in bacteria to achieve the native functional protein, including chaperone requirement and protease susceptibility. Future work will aim to better understand the interplay between clients, chaperones, and proteases to gain a broader view of the mechanisms of protein homeostasis in bacteria.

## Material and methods

### Strains, Plasmids and Growth conditions

Strains used in this study are listed in **Table S1**. The reference *S. oneidensis* strain used is MR1-R [42] and the reference strain for *E. coli* is MG1655 (K12) [43]. The strains *S. oneidensis* Δ*hsp90* and *E. coli* Δ*hsp90* were constructed previously [18,29]. *S. oneidensis and E. coli* WT or Δ*hsp90* strains with natural *tilS*_*So*_ or *tilS*_*Ec*_ replaced by *tilS*_*Ec*_ or *tilS*_*So*_, respectively, were constructed for this study as described in Supplementary Materials and Methods.

Plasmids used in this study are listed in **Table S2**. Plasmid construction is described in Supplementary Materials and Methods. Plasmids were introduced by conjugation in *S. oneidensis* and by transformation in *E. coli*.

Site-directed mutagenesis was performed with the Quickchange mutagenesis kit (Agilent) according to manufacturer’s instructions. Mutations were checked by sequencing. Sequences of the TilS_So_-TilS_Ec_ chimeras are indicated in **Table S3**. When necessary, chloramphenicol (25 µg/mL), kanamycin (50 µg/mL), ampicillin (50 µg/mL) or streptomycin (100 µg/mL) was added.

### Computational analysis of TilS orthologs

#### Taxonomic tree

To construct a taxonomic tree based on the TilS protein, we searched in InterPro IPR015262, which identifies the substrate-binding domain of TilS [44]. A total of approximately 18,500 proteins were retrieved, with the majority exhibiting two predominant domain architectures: IPR011063 -IPR015262 - IPR012796 (59%) and IPR011063 - IPR015262 (39%). Focusing on proteins possessing the IPR011063 - IPR015262 - IPR012796 architecture, we identified ∼10,500 proteins distributed across more than 6,000 bacterial species. Using the NCBI taxonomy [45], a taxonomic tree of these species was generated, demonstrating the widespread presence of TilS across bacteria. *Pseudomonadota* (Proteobacteria) were the most represented group, accounting for 72% of the sequences, followed by the *Bacillati* (Terrabacteria) group at 18%. The taxonomic analysis delves into the evolutionary distribution of TilS and its conservation across bacterial lineages. To further characterize TilS close to *S. oneidensis* and based on the phylogenetic analysis and the structure of TilS organization, we reduced the distribution of proteins to those best defined on Uniprot [26]. The reviewed TilS were recovered (with the addition of the TilS of *Shewanella xiamenensis*, due to its proximity to *S. oneidensis*, see **Table S4** for the sequences) and the C2 domains of these proteins were aligned with MUSCLE v3 [46]. Evolutionary history was inferred by using the Maximum Likelihood method and JTT matrix-based model with MEGA X [47]. The tree is drawn to scale, with branch lengths measured in the number of substitutions per site (**Table S5** for the Newick file). Based on this tree, we then selected the subtree presented (**Figure S6**), with the 73 sequences revolving around *E. coli* and *S. oneidensis* to select several sequences to perform experiments *in vivo*.

#### TilS sequence alignment

TilS_So_ and TilS_Sx_ sequences were aligned using “Multalin” tool (http://multalin.toulouse.inra.fr/multalin/).

The ESPript program (https://espript.ibcp.fr/ESPript/ESPript/) was used to display the alignment.

#### Thermal Shift Assays to determine TilS melting temperature

TilS proteins were purified as described in Supplementary Materials and Methods. 5 µg of proteins in 25 mM Tris-HCl pH 7.5, 50 mM KCl labeled with 10X SYPRO Orange (Sigma Life Science) were exposed to increasing temperatures from 15 to 90°C at a scan rate of 0.5°C per 30 s using BioRad CFX96 Touch RealTime PCR instrument. The protein unfolding curves were monitored by detecting changes in SYPRO Orange fluorescence. Melting temperatures were determined using the first derivative values of raw fluorescence data using Bio-Rad CFX Manager 3.1 software.

#### Microscale Thermophoresis

90 µL of TilS (340 nM) in PBS-Tween 0.05 %, pH 7.3, was labeled with 90 µL of 100 nM RED-tris-NTA 2^nd^ generation dye (NanoTemper). According to the manufacturer protocol, after 30 minutes incubation at room temperature, sample was centrifuged 10 minutes at 15 000 g and 10 µL of labeled TilS at 170 nM was transferred into 16 tubes containing 10 µL of serial 2-fold dilutions of Hsp90_So_ in PBS-Tween 0.05 %, pH 7.3. Final concentrations in capillary are 85 nM TilS and 50 µM Hsp90_So_ at highest concentration to 1.5 nM at the lowest. The resulting solutions were loaded onto 16 Monolith capillaries (NanoTemper). Fluorescence was measured at 670 nm at an ambient temperature of 25°C using Monolith NT.115 according to the manufacturer instructions. Instrument parameters were adjusted to 50% LED power and medium MST power. Data of two independently pipetted measurements were analyzed (MO.Affinity Analysis 3 software version 2.6.3, NanoTemper Technologies) using the signal from an MST on-time of 2.5 s.

#### Bacterial two-hybrid assays

Bacterial two-hybrid assays were performed as described by Battesti and Bouveret with modifications [28]. The Bth101 Δ*hsp90* strains lacking adenylate cyclase and Hsp90 corresponding genes were transformed with two plasmids. The first encoded the fusion protein between the T18 domain of *B. pertussis* adenylate cyclase at the N-terminal extremity with TilS from *E. coli, S. oneidensis* or *S. xiamenensis*, at the C-terminal extremity. The second encoded the fusion protein between the T25 domain of *B. pertussis* adenylate cyclase at the N-terminal extremity with Hsp90 from *S. oneidensis* or *E. coli* at the C-terminal extremity. As negative controls, vectors producing the T25 or T18 domains were used. Transformants were incubated at 28°C. After 3 days, colonies were inoculated in LB medium supplemented with ampicillin, kanamycin and isopropylthio-β-galactoside. After overnight culture at 28°C, cells were lysed with 1mg/mL lysozyme and Popculture Reagent solution (Millipore) for 15 minutes. Then, Z buffer was added (100 mM phosphate buffer pH=7, 10 mM KCl, 1 mM MgSO_4_, 50 mM β-mercaptoethanol). Before measurement, 2.2 mM orthonitrophenyl-β-galactoside was added. β-galactosidase activity was measured via a modified Miller assay adapted for use in a Tecan Spark microplate reader as described previously [48].

#### Western blot analysis to evaluate TilS amount

TilS amount was determined by Western blots. The p*tilS*-6his plasmids carrying *tilS* genes from several species were introduced into *S. oneidensis* or *E. coli* WT or Δ*hsp90* strains. The strains were grown at 28°C overnight with chloramphenicol, diluted to OD_600_ = 0.1, and incubated at 28°C or 34°C. After 3 h, 0.02% arabinose was added and 2 h later the same amount of cells was collected. Pellets were resuspended in denaturing loading buffer and heat treated at 95°C. Proteins were separated by SDS-PAGE and transferred by Western blot. TilS was detected using anti-6his antibody (Thermo). Hsp90 was detected with anti-Hsp90_So_ or anti-Hsp90_Ec_ antibody. As neutral loading control, a non-specific band detected by the anti-AtcJ antibody was used [48].

#### Bacterial growth of *S. oneidensis* or *E. coli*

These experiments were performed as previously described with modifications [20]. After overnight precultures at 28°C in LB medium, the *S. oneidensis* or *E. coli* strains were inoculated to OD_600_ = 0.1 in LB and incubated at 28°C until late exponential phase. For growth on liquid media, cells were diluted to OD_600_ = 0.0005 in LB. Growth was measured in a microplate reader at indicated temperatures.

## Supporting information

SI results, SI Materials and Methods, Tables S1 to S3, Figures S1 to S10

## CRediT authorship contribution statement

**Marie Corteggiani:** Conceptualization, Investigation, Validation, Visualization, Writing – original draft, Writing – review and editing. **Amine Ali-Chaouche:** Investigation, Validation, Writing – review and editing. **Miha Bahun:** Investigation, Validation, Writing – review and editing. **Flora Honoré:** Investigation, Validation, Writing – review and editing. **Deborah Byrne:** Resources, Validation, Writing – review and editing. **Sébastien Dementin:** Investigation, Validation, Writing – review and editing. **Mathieu E. Rebeaud:** Conceptualization, Investigation, Validation, Visualization, Writing – original draft, Writing – review and editing. **Olivier Genest:** Conceptualization, Investigation, Validation, Visualization, Writing – original draft, Writing – review and editing, Supervision, Project administration, Funding acquisition.

## Acknowledgement

We thank members of our groups for help and fruitful discussions, and Yann Denis from the transcriptomic platform of the IMM (CNRS) for assistance with thermal shift assay. M.E.R. would like to thank Paolo De Los Rios for helpful discussions and the opportunity to work on several interesting side projects. This work was supported by the Centre National de la Recherche Scientifique, Aix Marseille Université, and the Agence Nationale de la Recherche (ANR-16-CE11-0002-01 and ANR-20-CE44-0017).

