## Supplementary material for "Subtle variations in a client protein determine bacterial Hsp90 dependence": SI results, SI Materials and Methods, Tables S1 to S3, Figures S1 to S10

[a]Aix-Marseille Univ, CNRS, BIP UMR 7281, IMM, 31 Chemin Joseph Aiguier, 13402 Marseille, France.

[b]Department of Food Science and Technology, Biotechnical Faculty, University of Ljubljana, Jamnikarjeva 101, SI-1000, Ljubljana, Slovenia.

[c]Aix-Marseille Univ, CNRS, IMM-FR3479, 31 Chemin Joseph Aiguier, 13402, Marseille, France.

[d]Institute of Physics, School of Basic Sciences, École Polytechnique Fédérale de Lausanne - EPFL, Lausanne, Switzerland.

**This supplementary document includes:**

- Supplementary Results
- Supplementary Materials and Methods
- Supplementary Tables S1 to S3
- Supplementary References
- Supplementary Figures 1 to 10, and Legends

**Supplementary Results**

**Hsp90 dependence for TilS protection is not an evolutionary conserved process.**

Since $TilS_{Ec}$ is not dependent on Hsp90, but $TilS_{So}$ is, we hypothesized that we could link the evolution of TilS to its dependence on Hsp90. Based on TilS sequences, we built a phylogenetic tree with the sequence of the C2 domains of several TilS orthologs, including $TilS_{Ec}$ and $TilS_{So}$ (**Figure S6**). We expected that the TilS with C2 domains close to $TilS_{So}$ would be degraded in *S. oneidensis* Δ*hsp90*, whereas TilS with C2 sequences close to $TilS_{Ec}$ would be stable even in the absence of Hsp90. The C2 of $TilS_{So}$ is in the same subgroup as two other *Shewanella* species: *S. xiamenensis* and *S. woodyi*. The C2 of $TilS_{Ec}$ and TilS from *Yersinia pestis*, *Salmonella enterica* serovar Typhymurium, and *Vibrio cholera* also forms a subgroup. TilS of *Francisella tularensis* belongs to another group, distinct from the $TilS_{So}$ and $TilS_{Ec}$ subgroups.

We cloned several *tilS* genes representative of the different groups and subgroups of orthologous TilS proteins classified by their C2 including *Francisella tularensis*, *Shewanella woodyi*, *Shewanella xiamenensis*, *Vibrio cholera*, and *Salmonella enterica* serovar Typhymurium. The resulting plasmids were introduced into *S. oneidensis* WT or Δ*hsp90* strains, and the resulting strains were grown under heat stress (34°C) or permissive (28°C) temperatures. The amount of TilS was determined by Western blot (**Figure S7 A-B**) and quantification of the bands (**Figure S7 C-D**).

As expected, the TilS of *V. cholerae* and *Y. pestis* belonging to the $TilS_{Ec}$ subgroup were not degraded in the Δ*hsp90* strain at both temperatures suggesting that these two proteins are not dependent on the chaperone Hsp90. Surprisingly, TilS of *S. enterica* is less abundant in the absence of the chaperone both at 34°C and 28°C.

In contrast to our expectations, TilS of *S. xiamenensis* did not seem to be degraded in the absence of Hsp90. For *S. woodyi*, the variability between replicates did not allow us to draw a clear conclusion on the dependence of its TilS on Hsp90.

The TilS of *F. tularensis* which belongs to another C2 domain classification group, apart from $TilS_{Ec}$ and $TilS_{So}$, seems to be dependent on the chaperone at 34°C but not at 28°C, similarly to $TilS_{So}$.

Altogether these results suggest that it is not possible to draw a correlation between the amino acid sequences of the C2 domain of TilS and the dependence on the chaperone Hsp90. Surprisingly, TilS of *S. xiamenensis* which is not degraded in the absence of the chaperone, shares 92% identity with $TilS_{So}$.

### Supplementary Materials and Methods

**Strain constructions.** The *S. oneidensis* WT or Δ*hsp90* strains with natural $tilS_{So}$ replaced by $tilS_{Ec}$ or $tilS_{So}$(L340P), were constructed using homologous recombination as previously described with some modifications [1]. For the replacement of $tilS_{So}$ by $tilS_{Ec}$ gene, $tilS_{Ec}$ was amplified by PCR from *E. coli* MG1655 chromosome and 500 bp long flanking sequences upstream and downstream $tilS_{So}$ locus were amplified from *S. oneidensis* MR1-R chromosome. For the $tilS_{So}$(L340P) mutation, two flanking sequences of 500 bp upstream and downstream of the codon of interest were amplified. The resulting constructions were cloned in the pKNG101 suicide vector using NEBuilder® HiFi DNA Assembly kit (NEB) according to the manufacturer protocol, and subsequently introduced in *E. coli* CC118λpir strain. The resulting plasmids were introduced in *S. oneidensis* by conjugation. After integration into the chromosome, plasmids were removed using sucrose. For mutations on the chromosome of *E. coli*, the $tilS_{So}$ gene with 600 bp flanking sequences from upstream and downstream $tilS_{Ec}$ was amplified by PCR and cloned in the pKO3 suicide vector using NEBuilder® HiFi DNA Assembly kit (NEB) according to the manufacturer protocol. The resulting plasmid was transformed in *E. coli* MG1655 strain. After integration into the chromosome at 42°C because of thermosensitive replication of the pKO3, the plasmid was removed using sucrose. All the constructions were first checked by PCR with specific primers and further confirmed by sequencing.

**Plasmid constructions.** The pBad33 plasmids carrying *tilS* genes were obtained by PCR amplification of the $tilS_{Ec}$ and/ or $tilS_{So}$ genes from the chromosomes of *E. coli* MG1655 and *S. oneidensis* MR1-R, respectively. For the chimeric constructions, the sequences amplified from $tilS_{Ec}$ or $tilS_{So}$ are indicated in **Table S3**. Orthologous *tilS* other than $tilS_{Ec}$ or $tilS_{So}$ were amplified from genes synthetized by the Genecust company. The resulting fragments were cloned into the pBad33 vector digested with XmaI and XbaI restriction enzymes using NEBuilder Assembly kit (NEB).

The pT18 plasmids carrying *tilS* genes were obtained as described above. The pT18 plasmid was digested with EcoRI and XhoI enzymes. The pET24b plasmids carrying *tilS* were obtained as described above. The pET24b plasmids were digested with NdeI and SacI enzymes.

The pBad33 plasmids carrying $hsp90_{So}$ or $hsp90_{Ec}$ under the control of their own promoter were obtained by the amplification of $hsp90_{So}$ or $hsp90_{Ec}$ including their respective 344 bp upstream the ATG codon. $hsp90_{So}$ was inserted into pBad33 digested with XmaI and XbaI enzymes and $hsp90_{Ec}$ was inserted into pBad33 digested with KpnI using NEBuilder Assembly kit (NEB). All plasmid constructions were verified by sequencing.

**Growth of *S. oneidensis* or *E. coli*.** These experiments were performed as previously described with modifications [2]. After overnight precultures at 28°C in LB medium, the *S. oneidensis* or *E. coli* strains were inoculated to $OD_{600}$ = 0.1 in LB and incubated at 28°C until late exponential phase. For strains carrying pBad33 plasmid, chloramphenicol was added. For growth on liquid media, cells were diluted to $OD_{600}$ = 0.0005 in LB. Growth was measured in a microplate reader at indicated temperatures. For growth on solid media, cells were diluted to $OD_{600}$ = 1 and 3 µL of 10x serial dilutions were spotted on LB agar plates. Plates were incubated at 28°C overnight and at 35°C for 24 h.

**Proteins.** To purify $TilS_{So}$, $TilS_{Ec}$, $TilS_{Sx}$ and $TilS_{So}$L340P, the corresponding pET24b-*tilS* plasmids were introduced into the BL21 (DE3) strain by transformation. The resulting strains were grown in LB at 37°C. At $OD_{600}$ = 0.8, 0.1 mM isopropyl β-D-1-thiogalactopyranoside (IPTG) was added, and the strains were grown overnight at 16°C, cells were collected by centrifugation, lysed by French Press and centrifuged at 13 000 rpm, 4°C, 15 min. After a second centrifugation step at 45,000 rpm, 4°C, 45 min, the supernatant was loaded onto a HisTrap HP 1mL column (Cytiva), and purification was performed as described by the manufacturer. To improve purification, TilS extract from HisTrap column were loaded onto Gel Filtration Superdex® 16 600 Hiload S200 (Cytiva). $Hsp90_{So}$ purification was performed as described previously [2,3]. Purified proteins were stored at -80°C in 50 mM Tris-HCl pH=7.5, 100 mM KCl, 1 mM DTT, and 10% glycerol. Concentrations are given as monomeric proteins.

**Gel Filtration analysis.** 8 µM of proteins in 300 µL of 25mM Tris HCl pH 7.5, 50 mM KCl, 1 mM DTT and 10% glycerol were loaded onto gel filtration Superdex® 200 Increase 10 300 GL column (Cytiva).

**Limited proteolysis experiments.** $TilS_{So}$, $TilS_{Ec}$, $TilS_{Sx}$ or $TilS_{So}$L340P (15 µg) was mixed (50 µL final volume) with trypsin (0.25 µg) in 50 mM Tris-HCl buffer pH 7.5 containing 100 mM KCl, 10% glycerol and 1 mM DTT and incubated at 37°C. At the indicated times, 10 µL was removed, mixed with 5 µL of loading buffer, denatured for 5 min at 95°C and analyzed by SDS-PAGE. Trypsin was purchased from Sigma.

### Supplementary Tables

### Table S1: Strains used in this study

| STRAINS | CHARACTERISTICS | SOURCE |
|---|---|---|
| ***S. oneidensis* strains** | | |
| MR1-R | *S. oneidensis* MR1 strain ATCC 700550, Rifampicine resistance. | [5] |
| MR1-R Δ*hsp90*$_{So}$ | MR1-R with deletion of *hsp90* gene. | [3] |
| MR1-R *tilS*$_{Ec}$ | MR1-R with substitution of the whole *tilS*$_{So}$ gene by *tilS*$_{Ec}$. | This study |
| MR1-R Δ*hsp90*$_{So}$ *tilS*$_{Ec}$ | MR1-R with substitution of the whole *tilS*$_{So}$ gene by *tilS*$_{Ec}$ in Δ*hsp90*$_{So}$ background. | This study |
| MR1-R *tilS*$_{So}$(L340P) | MR1-R with mutations in *tilS* gene leading to L340P substitutions in TilS$_{So}$. | This study |
| MR1-R Δ*hsp90*$_{So}$ *tilS*$_{So}$(L340P) | MR1-R with mutations in *tilS* gene leading to L340P substitutions in TilS$_{So}$, in Δ*hsp90*$_{So}$ background. | This study |
| MR1-R *tilS*$_{So}$(L340P-A344E) | MR1-R with mutations in *tilS* gene leading to L340P-A344E substitutions in TilS$_{So}$. | This study |
| MR1-R Δ*hsp90*$_{So}$ *tilS*$_{So}$(L340P-A344E) | MR1-R with mutations in *tilS* gene leading to L340P-A344E substitutions in TilS$_{So}$, in Δ*hsp90*$_{So}$ background. | This study |
| ***E. coli* strains** | | |
| Bth101 Δ*hsp90*$_{Ec}$ | cya-, deleted of *hsp90*. | [6] |
| MG1655 | MG1655 strain. | [7] |
| MG1655 Δ*hsp90*$_{Ec}$ | MG1655 strain with deletion of *hsp90* gene. | [8] |
| MG1655 | MG1655 with substitution of the whole *tilS*$_{Ec}$ gene by *tilS*$_{So}$. | This study |
| MG1655 Δ*hsp90*$_{Ec}$ | MG1655 with substitution of the whole *tilS*$_{Ec}$ gene by *tilS*$_{So}$ in Δ*hsp90*$_{So}$ background. | This study |
| Bl21(DE3) | Used for protein overproduction. Codes the T7 RNA polymerase. | Novagen |
| 1047 /pRK2013 | *E. coli* "helper" strain for conjugation. | [9] |
| CC118λpir | *λpir* lysogens of *E. coli* CC118. Used for pKNG101 plasmid replication. | [10] |

### Table S2: Plasmids used in this study

| PLASMIDS | CHARACTERISTICS | SOURCE |
|---|---|---|
| **Plasmids used for measuring TilS amount** | | |
| pBad33 | pBad33 plasmid. Arabinose inducible. Chloramphénicol resistance. | [11] |
| p*tilS*$_{So}$ | pBad33 plasmid carrying *tilS*$_{So}$ gene and a sequence coding six histidines (6His) tag. Production of a fusion TilS protein with 6His tag in N-terminus. | [3] |
| p*tilS*$_{Ec}$ | pBad33 plasmid carrying *tilS*$_{Ec}$ gene and a sequence coding six histidines (6His) tag. Production of a fusion TilS protein with 6His tag in N-terminus. | This study |
| p*tilS*$_{EEES}$ | pBad33 plasmid carrying a chimeric *tilS* gene and a sequence coding six histidines (6His) tag. Production of the TilS chimera EEES protein fused with 6His tag in N-terminus. | This study |
| p*tilS*$_{EESS}$ | pBad33 plasmid carrying a chimeric *tilS* gene and a sequence coding six histidines (6His) tag. Production of the TilS chimera EESS protein fused with 6His tag in N-terminus. | This study |
| p*tilS*$_{ESSS}$ | pBad33 plasmid carrying a chimeric *tilS* gene and a sequence coding six histidines (6His) tag. Production of the TilS chimera ESSS protein fused with 6His tag in N-terminus. | This study |
| p*tilS*$_{SSSE}$ | pBad33 plasmid carrying a chimeric *tilS* gene and a sequence coding six histidines (6His) tag. Production of the TilS chimera SSSE protein fused with 6His tag in N-terminus. | This study |
| p*tilS*$_{SSEE}$ | pBad33 plasmid carrying a chimeric *tilS* gene and a sequence coding six histidines (6His) tag. Production of the TilS chimera SSEE protein fused with 6His tag in N-terminus. | This study |
| p*tilS*$_{SEEE}$ | pBad33 plasmid carrying a chimeric *tilS* gene and a sequence coding six histidines (6His) tag. Production of the TilS chimera SEEE protein fused with 6His tag in N-terminus. | This study |
| p*tilS*$_{Ft}$ | pBad33 plasmid carrying *tilS*$_{Ft}$ gene of *Francisella tularensis* and a sequence coding six histidines (6His) tag. Production of a fusion TilS protein with 6His tag in N-terminus. | This study |
| p*tilS*$_{St}$ | pBad33 plasmid carrying *tilS*$_{St}$ gene of *Salmonella enterica* serovar Thyphymurium and a sequence coding six histidines (6His) tag. Production of a fusion TilS protein with 6His tag in N-terminus. | This study |
| p*tilS*$_{Sw}$ | pBad33 plasmid carrying *tilS*$_{Sw}$ gene of *Shewanella woodyi* and a sequence coding six histidines (6His) tag. Production of a fusion TilS protein with 6His tag in N-terminus. | This study |

| | | |
|---|---|---|
| p*tilS*$_{Sx}$ | pBad33 plasmid carrying *tilS*$_{Sx}$ gene of *Shewanella xiamenensis* and a sequence coding six histidines (6His) tag. Production of a fusion TilS protein with 6His tag in N-terminus. | This study |
| p*tilS*$_{Yp}$ | pBad33 plasmid carrying *tilS*$_{Yp}$ gene of *Yersinia pestis* and a sequence coding six histidines (6His) tag. Production of a fusion TilS protein with 6His tag in N-terminus. | This study |
| p*tilS*$_{Vc}$ | pBad33 plasmid carrying *tilS*$_{Vc}$ gene of *Vibrio cholerae* and a sequence coding six histidines (6His) tag. Production of a fusion TilS protein with 6His tag in N-terminus. | This study |
| p*tilS*$_{So}$(L340P) | pBad33 plasmid carrying *tilS*$_{So}$ mutant leading to L340P substitution in TilS$_{So}$ and a sequence coding six histidines (6His) tag. Production of a fusion TilS protein with 6His tag in N-terminus. | This study |
| p*tilS*$_{So}$(L340P-A344E) | pBad33 plasmid carrying *tilS*$_{So}$ mutant leading to L340P-A344E substitutions in TilS$_{So}$ and a sequence coding six histidines (6His) tag. Production of a fusion TilS protein with 6His tag in N-terminus. | This study |
| p*tilS*$_{So}$(L340S) | pBad33 plasmid carrying *tilS*$_{So}$ mutant leading to L340S substitution in TilS$_{So}$ and a sequence coding six histidines (6His) tag. Production of a fusion TilS protein with 6His tag in N-terminus. | This study |
| p*tilS*$_{So}$(L340G) | pBad33 plasmid carrying *tilS*$_{So}$ mutant leading to L340G substitution in TilS$_{So}$ and a sequence coding six histidines (6His) tag. Production of a fusion TilS protein with 6His tag in N-terminus. | This study |
| p*tilS*$_{So}$ (8-Ct$_{Sx}$) | pBad33 plasmid carrying *tilS*$_{So}$ mutant leading to replacement of 7 C-terminus last residues of TilS$_{So}$ by 8 C-terminus last residues of TilS$_{Sx}$ and a sequence coding six histidines (6His) tag. Production of a fusion TilS protein with 6His tag in N-terminus. | This study |
| **Plasmids used for two-hybrid experiments** | | |
| pT18 | pUT18C-linker vector. Carries the gene coding T18 adenylate cyclase domain of *B. pertussis*. IPTG inducible. Ampicillin resistance. | [12] |
| pT18-hsp90$_{So}$ | pUT18C-linker plasmid carrying hsp90So gene. | [3] |
| pT18-*tilS*$_{So}$ | pUT18C-linker plasmid carrying *tilS*$_{So}$ gene. | [3] |
| pT18-*tilS*$_{Ec}$ | pUT18C-linker plasmid carrying *tilS*$_{Ec}$ gene. | This study |
| pT18-*tilS*$_{Sx}$ | pUT18C-linker plasmid carrying *tilS*$_{Sx}$ gene. | This study |
| pT18-*tilS*$_{So}$(L340P) | pUT18C-linker plasmid carrying *tilS*$_{So}$ mutant leading to L340P substitution in TilS$_{So}$. | This study |
| pT18-*tilS*$_{So}$(L340P-A344E) | pUT18C-linker plasmid carrying *tilS*$_{So}$ mutant leading to L340P-A344E substitutions in TilS$_{So}$. | This study |
| pT18-*tilS*$_{So}$(L340S) | pUT18C-linker plasmid carrying *tilS*$_{So}$ mutant leading to L340S substitution in TilS$_{So}$. | This study |
| pT25 | pKT25-linker vector. Carries the gene coding T25 adenylate cyclase domain of *B.* | [12] |

| | | |
|---|---|---|
| | *pertussis*. IPTG inducible. Kanamycin resistance. | |
| pT25-*hsp90*$_{So}$ | pKT25-linker plasmid carrying *hsp90*$_{So}$ gene. | [3] |
| pT25-*hsp90*$_{Ec}$ | pKT25-linker plasmid carrying *hsp90*$_{Ec}$ gene. | [13] |
| **Plasmids for complementation experiments** | | |
| p*hsp90*$_{So}$ (own promoter) | pBad33 plasmid allowing the expression of *hsp90*$_{So}$ under the control of its own promoter. | This study |
| p*hsp90*$_{Ec}$ (own promoter) | pBad33 plasmid allowing the expression of *hsp90*$_{Ec}$ under the control of its own promoter. | This study |
| **Plasmids for protein purification** | | |
| pET24b-*tilS*$_{So}$ | Plasmid allowing the production of $TilS_{So}$ and a sequence coding six histidines (6His) tag. Production of a fusion TilS protein with 6His tag in C-terminus. | [3] |
| pET24b-*tilS*$_{Ec}$ | Plasmid allowing the production of $TilS_{Ec}$ and a sequence coding six histidines (6His) tag. Production of a fusion TilS protein with 6His tag in C-terminus. | This study |
| pET24b-*tilS*$_{Sx}$ | Plasmid allowing the production of $TilS_{Sx}$ and a sequence coding six histidines (6His) tag. Production of a fusion TilS protein with 6His tag in C-terminus. | This study |
| pET24b-*tilS*$_{So}$(L340P) | Plasmid allowing the production of $TilS_{So}$ L340P and a sequence coding six histidines (6His) tag. Production of a fusion TilS protein with 6His tag in C-terminus. | This study |
| pET24b-*hsp90*$_{So}$ | Plasmid allowing the production of $Hsp90_{So}$ wild-type. | [3] |
| **Suicide vectors** | | |
| pKNG101 | Suicide vector used for gene substitution and mutagenesis in *S. oneidensis*. Carries the *sacB* cassette which causes sensitivity to sucrose, and a λ-pir-dependent origin of replication. Streptomycin resistance. | [14] |
| pKO3 | Suicide vector used for gene substitution and mutagenesis in *E. coli*. Carries the *sacB* cassette which causes sensitivity to sucrose, and a thermosensitive origin of replication. Chloramphenicol resistance. | [15] |

**Table S3: Sequences of the chimeras**

| |
|---|
| >TilS_Ecoli_Uniprot_P52097<br>MTLTLNRQLLTSRQILVAFSGGLDSTVLLHQLVQWRTENPGVALRAIHVHHGLSANADAWVTHCENVCQQWQVPLVVERVQLAQEGLG<br>IEAQARQARYQAFARTLLPGEVLVTAQHLDDQCETFLLALKRGSGPAGLSAMAEVSEFAGTRLIRPLLARTRGELVQWARQYDLRWIEDES<br>NQDDSYDRNFLRLRVVPLLQQRWPHFAEATARSAALCAEQESLLDELLADDLAHCQSPQGTLQIVPMLAMSDARRAAIIRRWLAGQNAP<br>MPSRDALVRIWQEVALAREDASPCLRLGAFEIRRYQSQLWWIKSVTGQSENIVPWQTWLQPLELPAGLGSVQLNAGGDIRPPRADEAVS<br>VRFKAPGLLHIVGRNGGRKLKKIWQELGVPPWLRDTTPLLFYGETLIAAAGVFVTQEGVAEGENGVSFVWQKTLS |
| >TilS_Soneidensis_Uniprot_Q8EGF9<br>MTAQDLSVHIARWLDSLPLQAGSKLVLAYSGGVDSEVLAYGLSEYAKQRPDLRYQLIYVHHGLSPNADNWAKHCQARAAIYGLPVTVERV<br>QLILGPRVSVEAEARKARYQAILPHLNPQDILLTAHHEDDQLETILLALKRGQGPKGLAAMGQIQPLSLADKGSCLQVRPLLDISREMIETFA<br>QTRQLVHIEDESNQDDKYDRNFLRLEIIPRLKARWPSIATTASRSAQLCAEQQAIVETEVSERLPKLLVKAPVTEQTVLKLSELAAQPIEWQGI<br>LLRGFIESQEFSLPSYVQLQQMLQQLIHAKEDAKVHIRINDCVLRRFAGMLYLDSGETLSTALHITARDLHQEILTLLTQASAMVEDKIVPFAL<br>VTTGPRLRLPKADEVVSLGYGLPGQFRCQPHFRDKGRELKKLWQECAVPPWLRAEVGFLFYNDKLVMAFGLWVEKAFCAQGDEIGLSYLI<br>ANP |
| Chimera 1 : EEES<br>MTLTLNRQLLTSRQILVAFSGGLDSTVLLHQLVQWRTENPGVALRAIHVHHGLSANADAWVTHCENVCQQWQVPLVVERVQLAQEGLG<br>IEAQARQARYQAFARTLLPGEVLVTAQHLDDQCETFLLALKRGSGPAGLSAMAEVSEFAGTRLIRPLLARTRGELVQWARQYDLRWIEDES<br>NQDDSYDRNFLRLRVVPLLQQRWPHFAEATARSAALCAEQESLLDELLADDLAHCQSPQGTLQIVPMLAMSDARRAAIIRRWLAGQNAP<br>MPSRDALVRIWQEVALAREDASPCLRLGAFEIRRYQSQLWWIKSVGETLSTALHITARDLHQEILTLLTQASAMVEDKIVPFALVTTGPRLRL<br>PKADEVVSLGYGLPGQFRCQPHFRDKGRELKKLWQECAVPPWLRAEVGFLFYNDKLVMAFGLWVEKAFCAQGDEIGLSYLIANP |
| Chimera 2 : EESS<br>MTLTLNRQLLTSRQILVAFSGGLDSTVLLHQLVQWRTENPGVALRAIHVHHGLSANADAWVTHCENVCQQWQVPLVVERVQLAQEGLG<br>IEAQARQARYQAFARTLLPGEVLVTAQHLDDQCETFLLALKRGSGPAGLSAMAEVSEFAGTRLIRPLLARTRGELVQWARQYDLRWIEDES<br>NQDDSYDRNFLRLRVVPLLQQRWPHFAEATARSAALCAEQESLLDELLADDLAHCQSLVKAPVTEQTVLKLSELAAQPIEWQGILLRGFIES<br>QEFSLPSYVQLQQMLQQLIHAKEDAKVHIRINDCVLRRFAGMLYLDSGETLSTALHITARDLHQEILTLLTQASAMVEDKIVPFALVTTGPRL<br>RLPKADEVVSLGYGLPGQFRCQPHFRDKGRELKKLWQECAVPPWLRAEVGFLFYNDKLVMAFGLWVEKAFCAQGDEIGLSYLIANP |
| Chimera 3 : ESSS<br>MTLTLNRQLLTSRQILVAFSGGLDSTVLLHQLVQWRTENPGVALRAIHVHHGLSANADAWVTHCENVCQQWQVPLVVERVQLAQEGLG<br>IEAQARQARYQAFARTLLPGEVLVTAQHLDDQCETFLLALKRGSGPAGLSAMAEVSEFAGTRLIRPLLARTRGELVQWARQYDLRWIEDES<br>NQDDSYDRNFLRLRVVPLLQQRWPSIATTASRSAQLCAEQQAIVETEVSERLPKLLVKAPVTEQTVLKLSELAAQPIEWQGILLRGFIESQEFS<br>LPSYVQLQQMLQQLIHAKEDAKVHIRINDCVLRRFAGMLYLDSGETLSTALHITARDLHQEILTLLTQASAMVEDKIVPFALVTTGPRLRLPK<br>ADEVVSLGYGLPGQFRCQPHFRDKGRELKKLWQECAVPPWLRAEVGFLFYNDKLVMAFGLWVEKAFCAQGDEIGLSYLIANP |
| Chimera 4 : SSSE<br>MTAQDLSVHIARWLDSLPLQAGSKLVLAYSGGVDSEVLAYGLSEYAKQRPDLRYQLIYVHHGLSPNADNWAKHCQARAAIYGLPVTVERV<br>QLILGPRVSVEAEARKARYQAILPHLNPQDILLTAHHEDDQLETILLALKRGQGPKGLAAMGQIQPLSLADKGSCLQVRPLLDISREMIETFA<br>QTRQLVHIEDESNQDDKYDRNFLRLEIIPRLKARWPSIATTASRSAQLCAEQQAIVETEVSERLPKLLVKAPVTEQTVLKLSELAAQPIEWQGI<br>LLRGFIESQEFSLPSYVQLQQMLQQLIHAKEDAKVHIRINDCVLRRFAGMLYLDSTGQSENIVPWQTWLQPLELPAGLGSVQLNAGGDIRP<br>PRADEAVSVRFKAPGLLHIVGRNGGRKLKKIWQELGVPPWLRDTTPLLFYGETLIAAAGVFVTQEGVAEGENGVSFVWQKTLS |
| Chimera 5 : SSEE<br>MTAQDLSVHIARWLDSLPLQAGSKLVLAYSGGVDSEVLAYGLSEYAKQRPDLRYQLIYVHHGLSPNADNWAKHCQARAAIYGLPVTVERV<br>QLILGPRVSVEAEARKARYQAILPHLNPQDILLTAHHEDDQLETILLALKRGQGPKGLAAMGQIQPLSLADKGSCLQVRPLLDISREMIETFA<br>QTRQLVHIEDESNQDDKYDRNFLRLEIIPRLKARWPSIATTASRSAQLCAEQQAIVETEVSERLPKLPQGTLQIVPMLAMSDARRAAIIRRWL<br>AGQNAPMPSRDALVRIWQEVALAREDASPCLRLGAFEIRRYQSQLWWIKSVTGQSENIVPWQTWLQPLELPAGLGSVQLNAGGDIRPPR<br>ADEAVSVRFKAPGLLHIVGRNGGRKLKKIWQELGVPPWLRDTTPLLFYGETLIAAAGVFVTQEGVAEGENGVSFVWQKTLS |
| Chimera 6 : SEEE<br>MTAQDLSVHIARWLDSLPLQAGSKLVLAYSGGVDSEVLAYGLSEYAKQRPDLRYQLIYVHHGLSPNADNWAKHCQARAAIYGLPVTVERV<br>QLILGPRVSVEAEARKARYQAILPHLNPQDILLTAHHEDDQLETILLALKRGQGPKGLAAMGQIQPLSLADKGSCLQVRPLLDISREMIETFA<br>QTRQLVHIEDESNQDDKYDRNFLRLEIIPRLKARWPHFAEATARSAALCAEQESLLDELLADDLAHCQSPQGTLQIVPMLAMSDARRAAIIR<br>RWLAGQNAPMPSRDALVRIWQEVALAREDASPCLRLGAFEIRRYQSQLWWIKSVTGQSENIVPWQTWLQPLELPAGLGSVQLNAGGDI<br>RPPRADEAVSVRFKAPGLLHIVGRNGGRKLKKIWQELGVPPWLRDTTPLLFYGETLIAAAGVFVTQEGVAEGENGVSFVWQKTLS |

### Supplementary Figures

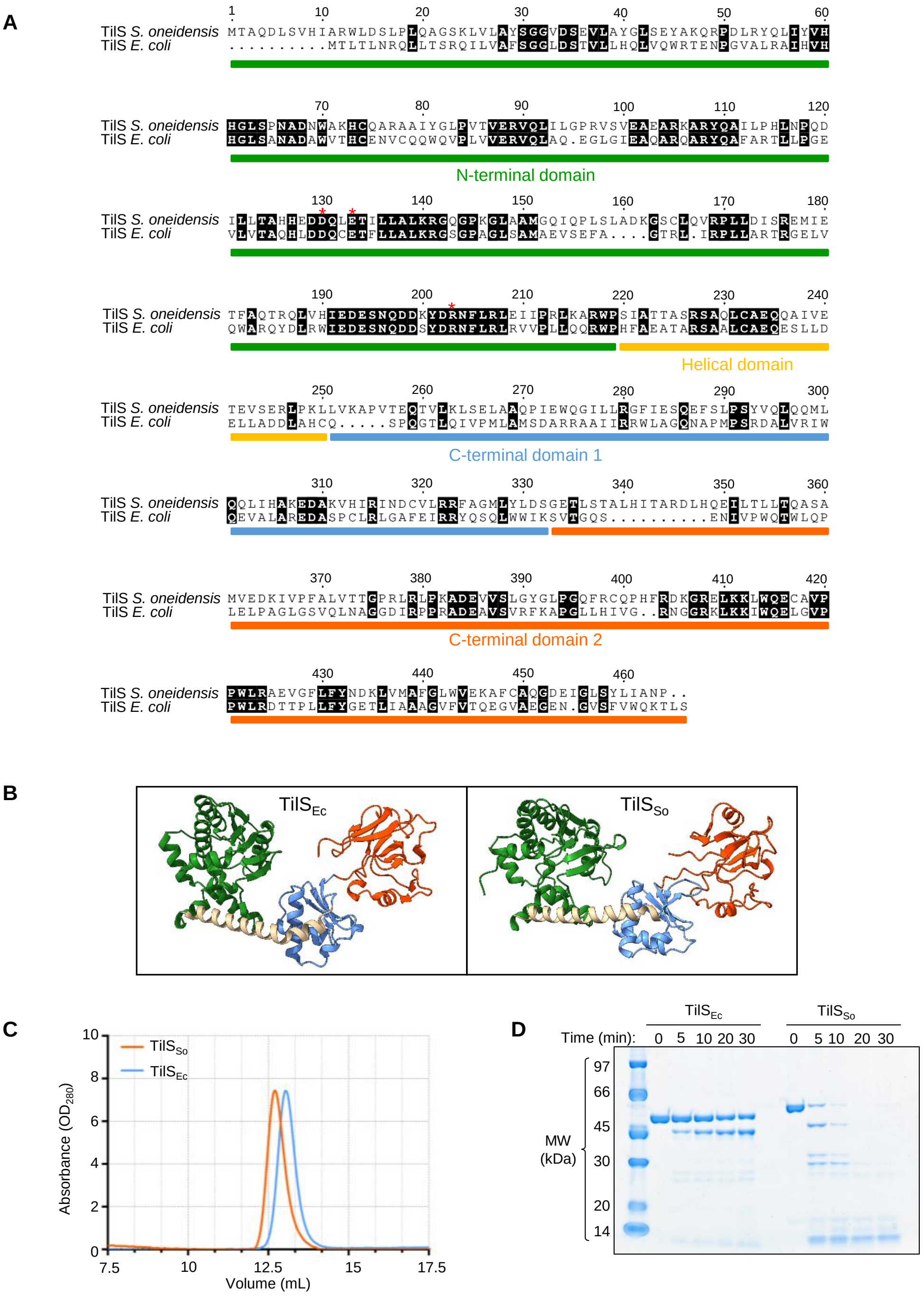


**Figure S1: A comparison between TilS of *E. coli* (TilS$_{Ec}$) and TilS of *S. oneidensis* (TilS$_{So}$).** (A) Sequence alignments of TilS$_{So}$ and TilS$_{Ec}$ using the multiple sequence alignment tool «Multalin» (http://multalin.toulouse.inra.fr/multalin/). The ESPript program (https://espript.ibcp.fr/ESPript/ESPript/) was used to display the alignment. The four domains of TilS$_{So}$ are represented: the N-terminal domain (green), the Helical domain (wheat), the C-terminal 1 domain (blue) and the C-terminal 2 domain (orange). The key catalytic residues are indicated by red asterisk. (B) Structure of TilS$_{Ec}$ (PDB 1NI5) and model of TilS$_{So}$ generated with AlphaFold 3.0 Software, pTM = 0.81. Each TilS domain is colored as in *A*. (C) Gel filtration analysis. Purified TilS$_{So}$ or TilS$_{Ec}$ (8 µM, 100 µL) were analyzed on a Superdex 200 Increase 10/300 GL size exclusion chromatography column. Normalized values of absorbance at 280 nm are shown. (D) Limited proteolysis profiles of purified TilS$_{Ec}$ and TilS$_{So}$ (15 µg) digested with trypsin (0.25 µg) for various times at 37°C in Tris-HCl buffer (50 mM, pH 7.5) containing 100 mM KCl, 10% glycerol and 1 mM DTT in a final reaction volume of 50 µL. At the indicated times, 10 µL was removed and analyzed by SDS-PAGE.

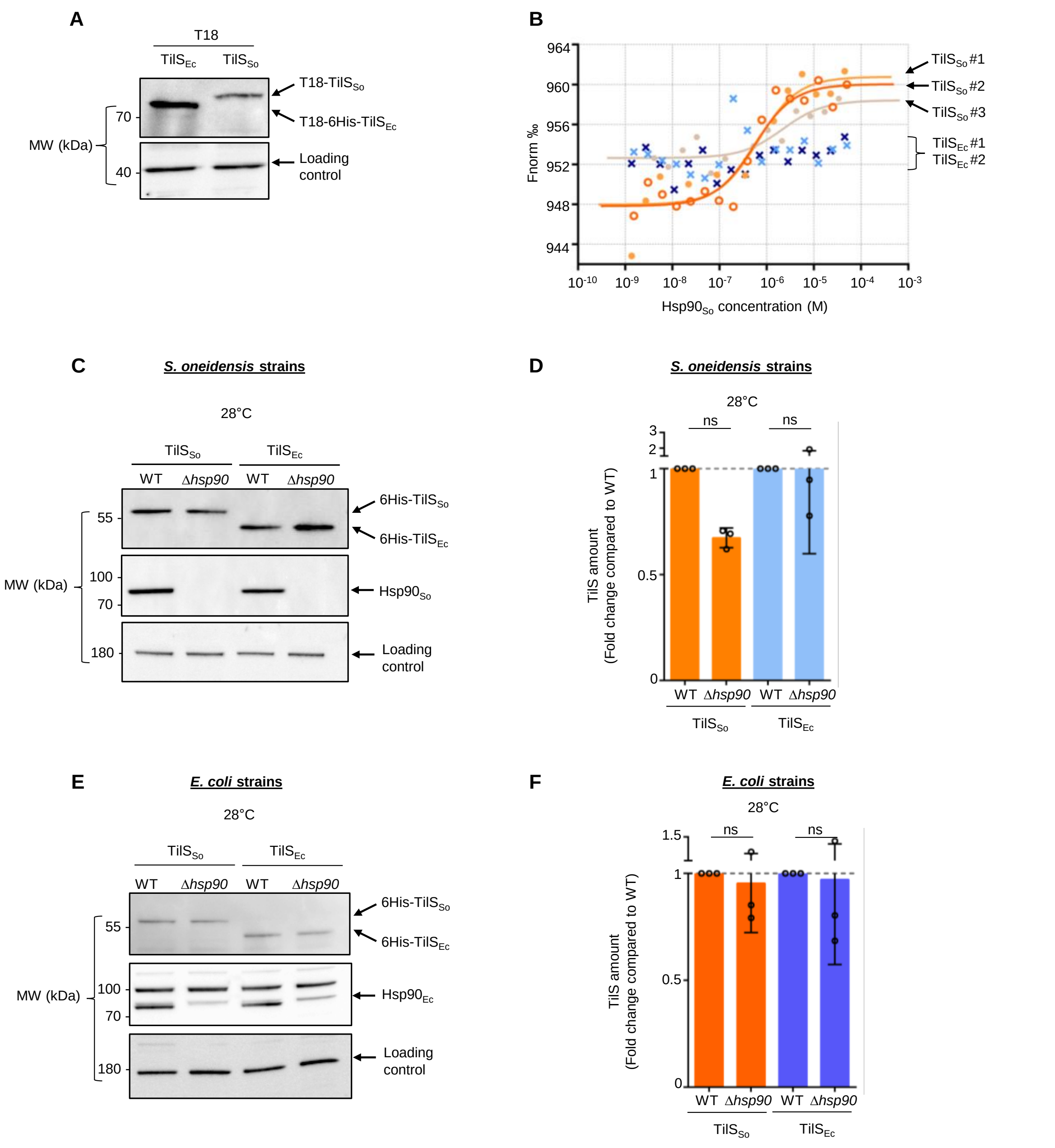


**Figure S2: Controls related to Figure 2.** (A) Western blots showing the production of T18-TilS$_{Ec}$ and T18-TilS$_{So}$ used in bacterial two-hybrid experiments. Anti-RecA antibody was used as neutral loading control. *E. coli* strains with plasmid allowing the production of T18-TilS$_{Ec}$ or TilS$_{So}$ were grown at 28°C overnight with 1mM IPTG to allow the expression of the corresponding genes. After 12 hours, samples were analyzed by Western blot using anti-Adenylate cyclase (T18) antibody. (B) Replicate of microscale thermophoresis experiment to determine the interaction between Hsp90 and TilS$_{So}$ or TilS$_{Ec}$. 85 nM of TilS labeled with RED-tris-NTA 2nd generation dye (NanoTemper) was added to serial 2-times dilutions of Hsp90$_{So}$ up to 50 µM. Fluorescence at 670 nm was measured, and values at 2.5 s were used to determine $K_d$. (C) Western blots showing the amount of TilS$_{So}$ or TilS$_{Ec}$ in *S. oneidensis*. The strains WT or *Δhsp90* with plasmid allowing the production of TilS$_{So}$ or TilS$_{Ec}$ with hexahistidine tag were grown at 28°C and 0.02% arabinose was added. Two hours later, samples were analyzed by Western blot using anti-His antibody. Anti-Hsp90$_{So}$ antibody was used as a control and a contaminant band revealed by the anti-AtcJ antibody was used as neutral sample control. (D) Quantification of anti-His Western blot shown in C. The peaks corresponding to the pixels of each band were quantified using ImageJ software. The amount of TilS$_{So}$ or TilS$_{Ec}$ in WT strain was set to 1, and each WT is compared to *Δhsp90* producing the corresponding TilS$_{So}$ or TilS$_{Ec}$ respectively. Data from three replicates are shown as mean ± SD. (E) Western blots showing the amount of TilS$_{So}$ or TilS$_{Ec}$ in *E. coli*. The protocol is similar to what was done in *C*. Anti-Hsp90$_{Ec}$ was used as control and a contaminant band revealed by the anti-AtcJ antibody was used as neutral loading control. (F) Quantification of anti-His Western blot shown in *E*. The method used is similar to *D*. In *A, C*, and *E*, the Western blots shown are representative of three independent experiments. In *D* and *F*, results of one-way ANOVA indicate whether the differences are significant or not (ns, $P > 0.05$).

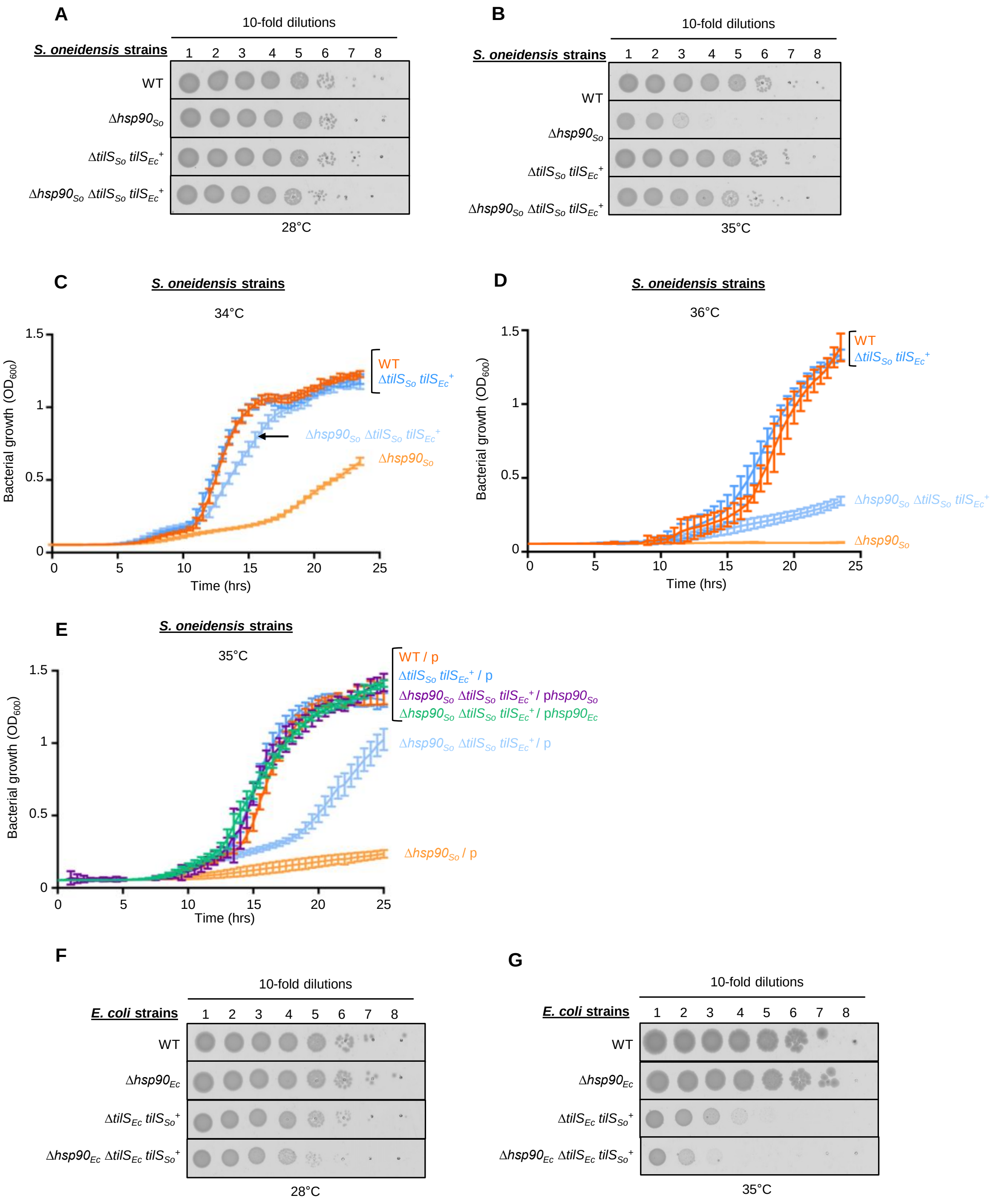


**Figure S3: Growth of the strains used in Fig. 3.** (A) Bacterial growth on solid media of *S. oneidensis* strains carrying *tilS* from *E. coli*. The strains *S. oneidensis* MR-1 wild-type (WT), deleted of *hsp90* (*Δhsp90*) with natural TilS or replaced by TilS from *E. coli* (Δ$tilS_{So}$ $tilS_{Ec}^{+}$) were grown at 28°C in liquid media and 10-fold serial diluted to be spotted on LB plates. The plates were incubated for 24 hours at 28°C. (B) Bacterial growth on solid media of the strains used in *A*, at 35°C. (C) Bacterial growth of *S. oneidensis* strains carrying *tilS* from *E. coli*. The strains *S. oneidensis* MR-1 wild-type (WT), deleted of *hsp90* (*Δhsp90*) with natural TilS or replaced by TilS from *E. coli* (Δ$tilS_{So}$ $tilS_{Ec}^{+}$) were grown in microplates in LB medium at 34°C with shaking for 24 hours. (D) Bacterial growth of *S. oneidensis* strains presented in *C*. The experiments is the same as in *C* except that strains were grown at 36°C. (E) Complementation of bacterial growth of *S. oneidensis* strains carrying *tilS* from *E. coli*. The strains *S. oneidensis* MR-1 wild-type (WT), deleted of *hsp90* (*Δhsp90*) with natural TilS or replaced by TilS from *E. coli* (Δ$tilS_{So}$ $tilS_{Ec}^{+}$) with empty vector or allowing the production of Hsp90 from *E. coli* ($hsp90_{Ec}$) or from *S. oneidensis* ($hsp90_{So}$) were grown in microplates in LB medium at 34°C with shaking for 24 hours. (F) Bacterial growth on solid media of *E. coli* strains carrying *tilS* from *S. oneidensis*. The strains *E. coli* MG1655 wild-type (WT), deleted of *hsp90* (*Δhsp90*) with natural TilS or replaced by

TilS from *S. oneidensis* ($\Delta tilS_{Ec}$ $tilS_{So}^{+}$) were grown at 28°C in liquid media and 10-fold serial diluted to be spotted on LB plates. The plates were incubated for 24 hours at 28°C. (G) Bacterial growth on solid media of the strains used in *E*, at 35°C. In *A, B, F* and *G*, the plates are representative of three independent experiments. In *C*, *D*, and *E*, data from at least three replicates are shown as mean ± SD.

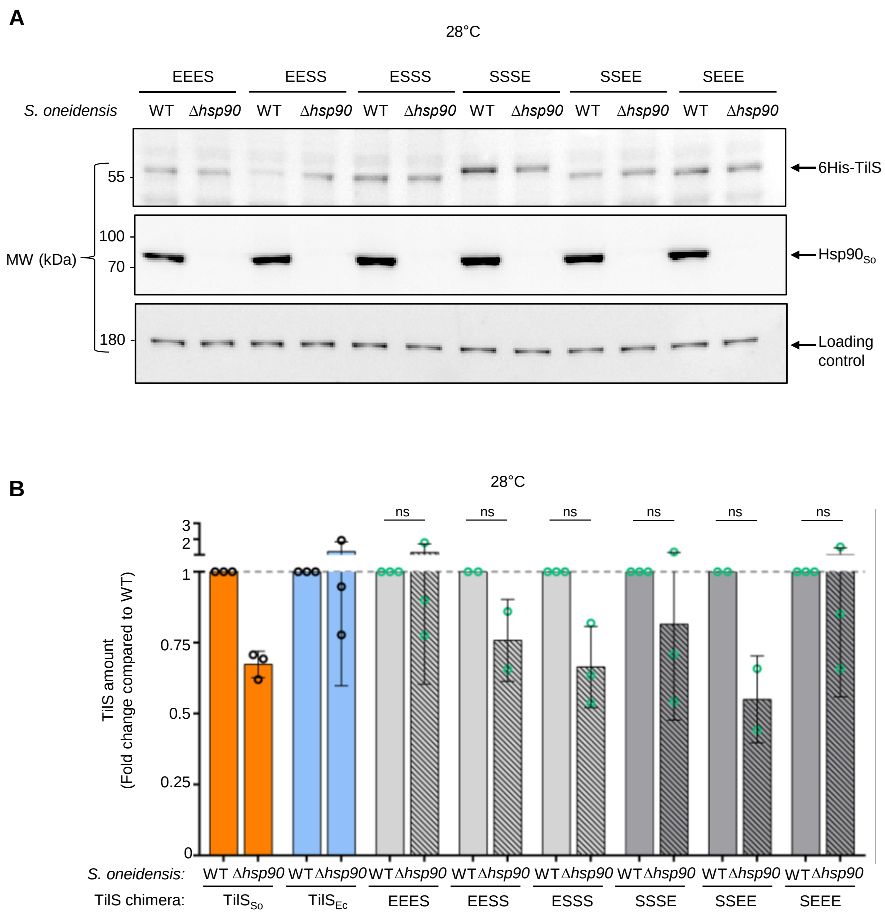


**Figure S4: TilS chimeras are not dependent on Hsp90 at 28°C.** (A) Western blot showing the abundance of TilS chimera in *S. oneidensis* WT or Δ*hsp90* at 28°C. The strains WT or Δ*hsp90* with plasmid pBad33 allowing the production of 6His-TilS chimera were grown at 28°C, and 0.02% arabinose was added. After 2 hours, samples were analyzed by Western blot with anti-His antibody. Anti-$Hsp90_{So}$ was used as a control and Anti-AtcJ as neutral loading control. This Western blot is representative of at least two independent experiments. (B) Quantification of the Western blot shown in *A*. The peaks corresponding to the pixels of each band were quantified using ImageJ software. The amount of each chimera in WT strain was set to 1, and each WT is compared to *Δhsp90* producing the corresponding chimera respectively. Data from two replicates are shown as mean ± SD. Results of one-way ANOVA indicate whether the differences are significant or not (ns, $P > 0.05$).

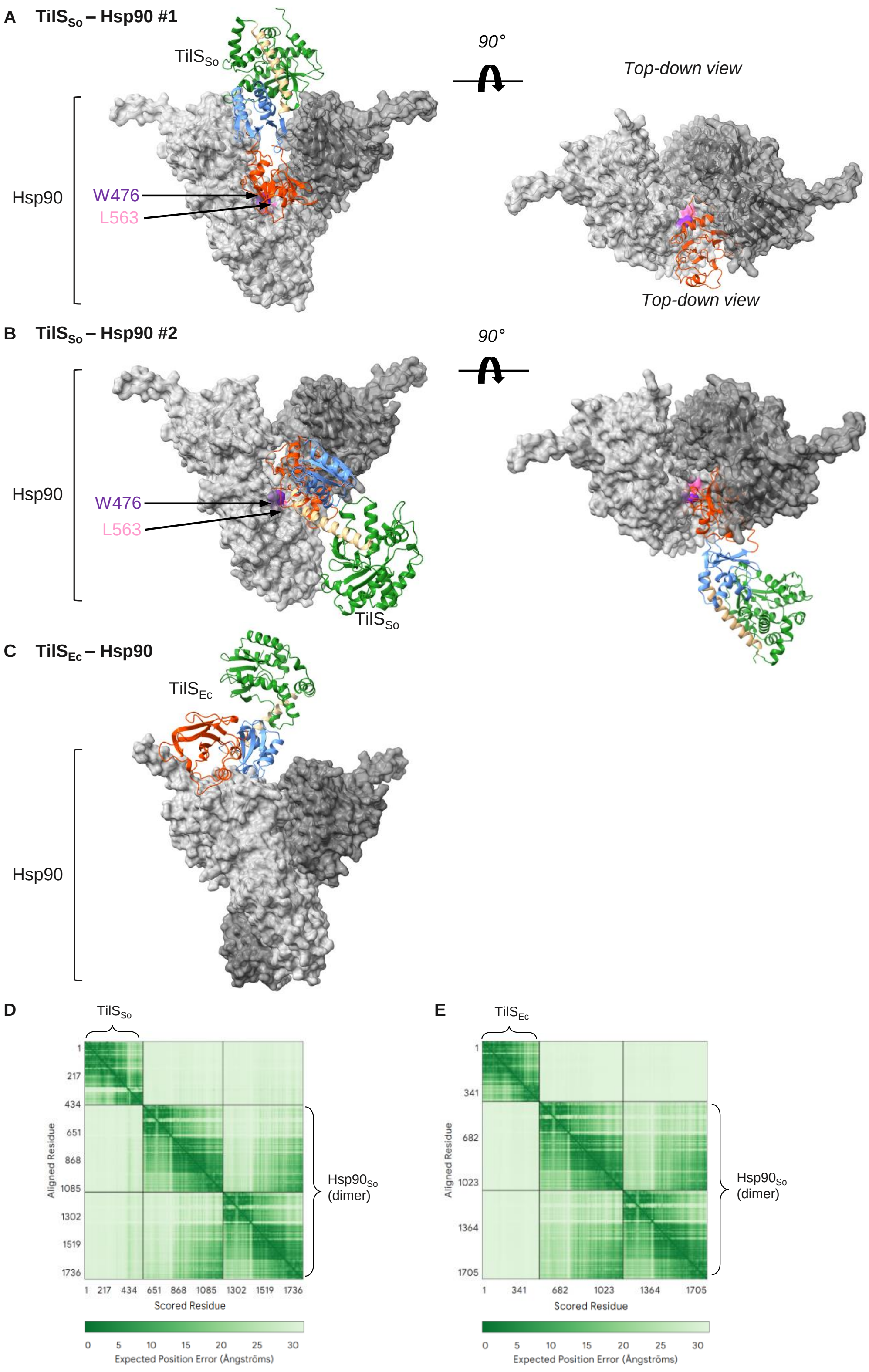


**Figure S5: AlphaFold 3 prediction models of the interaction of Hsp90 with $TilS_{So}$ or $TilS_{Ec}$.** (A-B) Prediction model of a monomer of $TilS_{So}$ (Uniprot : Q8EGF9) with a dimer of $Hsp90_{So}$ (Uniprot : Q8EFF7) by AlphaFold 3.0 software, ipTM = 0.38 pTM = 0.47. $TilS_{So}$ is colored by domains: the N-terminal domain (green), the Helical domain (wheat), the C-terminal 1 domain (blue) and the C-terminal 2 domain (orange). The W476 (purple) and L563 (pink) residues of Hsp90 important for the interaction with clients are shown. Among the five models presented by the software, four are similar to the model presented in *A*, and one shows $TilS_{So}$ as in *B*. In *A* right panel, $TilS_{So}$ structure was truncated to better visualize the insertion of the C-terminal 2 domain into the cleft formed by $Hsp90_{So}$. (C) Prediction model of a monomer of $TilS_{Ec}$ (Uniprot : P52097 )

with a dimer of $Hsp90_{So}$ (Uniprot : Q8EFF7) by AlphaFold 3.0 software, ipTM = 0.39 pTM = 0.5. $TilS_{Ec}$ is colored by domains as in *A*. (D-E) Heatmaps showing the predicted interaction confidence between $TilS_{So}$ (D) or $TilS_{Ec}$ (E) and the $Hsp90_{So}$ dimer, based on residue-residue contact probabilities.

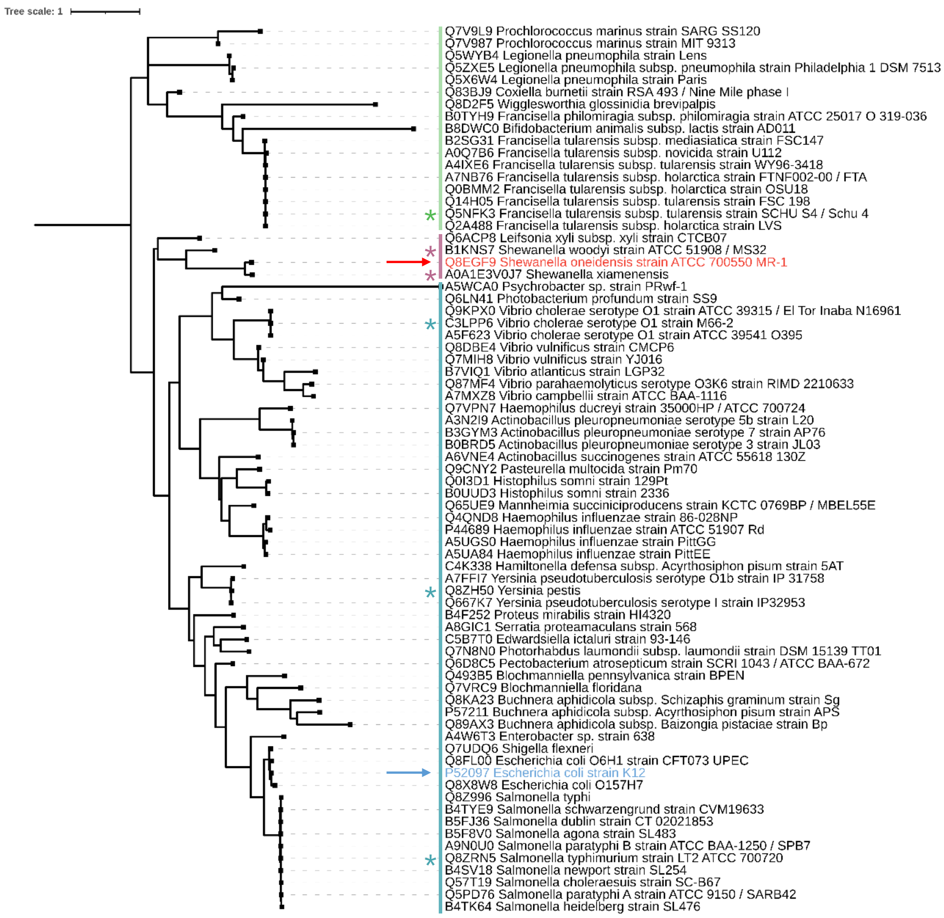


**Figure S6: Phylogenetic tree of the TilS C2 domain in selected bacterial species.** This maximum-likelihood phylogenetic tree illustrates the evolutionary relationships of the TilS C2 domain among bacterial taxa. Branches are labelled with UniProt ID and species names. Selected taxa are *Shewanella oneidensis* MR-1 (red) and *Escherichia coli* K12 (blue). Branches length indicate the substitution rate per site. Sequences of interest are indicated by asterisks.

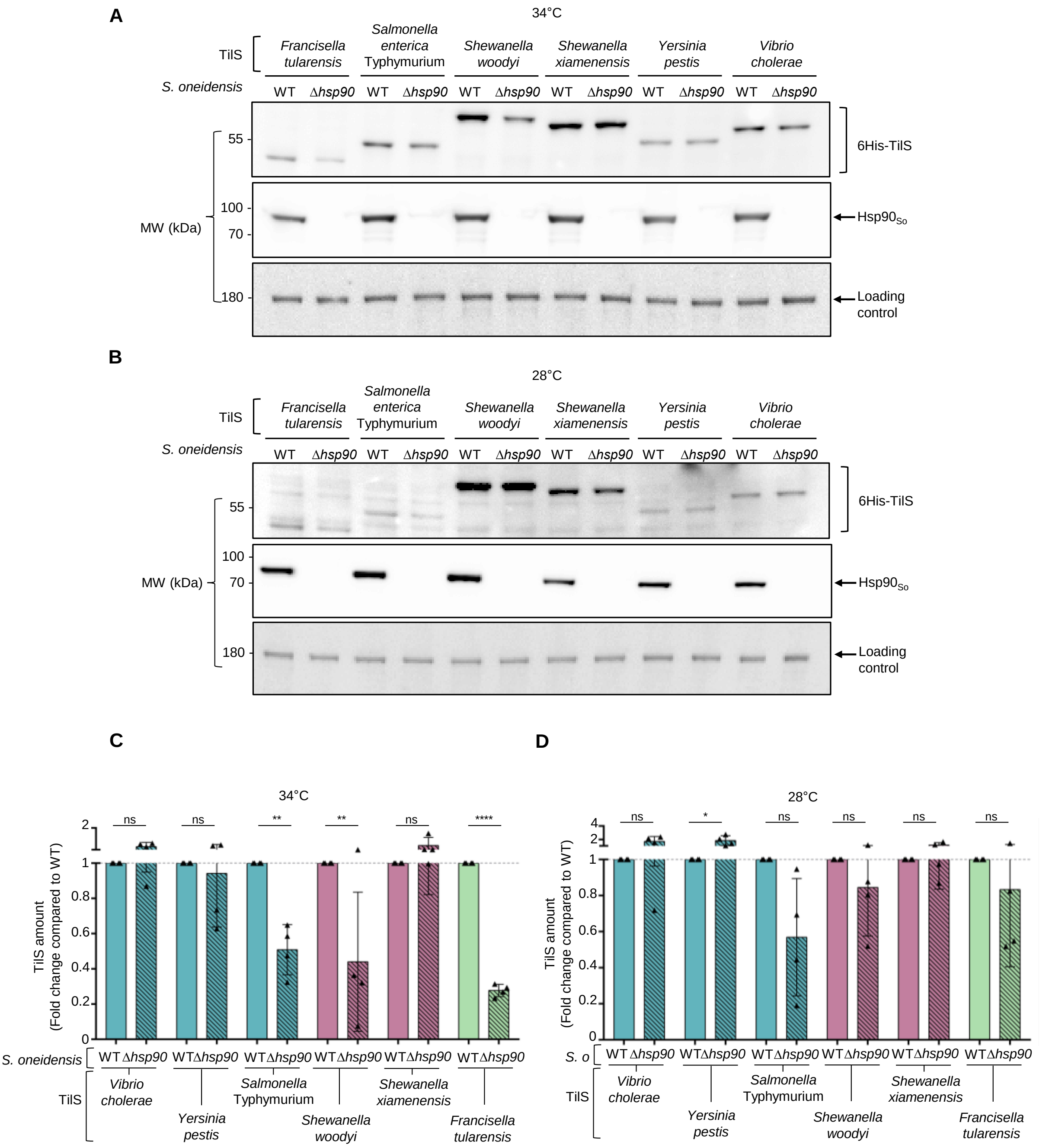


**Figure S7: Controls related to TilS orthologues.** (A) Western blot showing the amount of TilS orthologues. The strains of *S. oneidensis* WT or *Δhsp90* with plasmid allowing the production of TilS from *Shewanella woodyi*, *Shewanella xiamenensis*, *Vibrio cholerae*, *Francisella tularensis*, *Yersinia pestis* or *Salmonella enterica* serovar Typhymurium with hexahistidine tag were grown at 34°C (heat stress conditions) and 0.02% arabinose was added. Two hours later, samples were analyzed by Western blot using anti-His antibody. Anti-$Hsp90_{So}$ antibody was used as a control and anti-AtcJ antibody was used as neutral sample control. (B) Same experiment as in *B*, except that the strains were grown at 28°C. (C-D) Quantification of the Western blot shown in *A and B*. The peaks corresponding to the pixels of each band were quantified using ImageJ software. The amount of each chimera in WT strain was set to 1, and each WT is compared to *Δhsp90* producing the corresponding chimera respectively. Data from four replicates are shown as mean ± SD. Results of one-way ANOVA indicate whether the differences are significant (**P* ≤ 0.05, ***P* ≤ 0.01, *****P* ≤ 0.0001) or not (ns, *P* > 0.05).

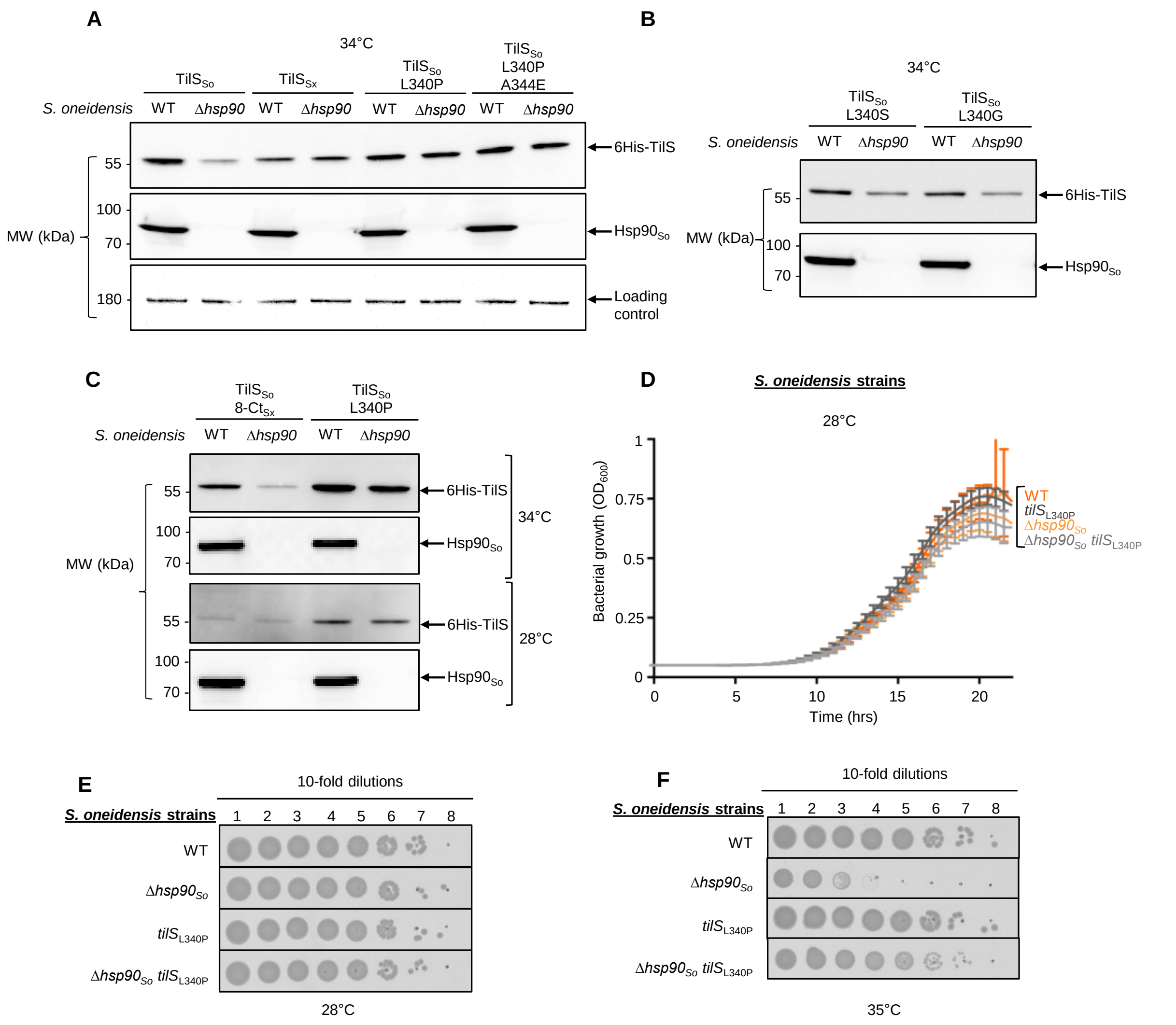


**Figure S8: Effects of a single mutation on TilS$_{So}$.** (A) Western blot showing the amount of TilS$_{So}$ WT or mutants or TilS$_{Sx}$ in *S. oneidensis* WT or *Δhsp90* strains. The strains WT or *Δhsp90* with plasmid allowing the production of TilS$_{So}$ WT or mutants or TilS$_{Sx}$ with hexahistidine tag were grown at 34°C and 0.02% arabinose was added. Two hours later, samples were analyzed by Western blot using anti-His antibody. As a control, anti-Hsp90 antibody was used. The Western blot is representative of three independent experiments. (B) Western blot showing the amount of TilS$_{So}$ L340S and L340G mutants in *S. oneidensis* WT or *Δhsp90*. The experiment was performed as described in *A*. (C) Western blot showing the amount of TilS$_{So}$ with the 7 last residues replaced by the 8 last residues of TilS$_{Sx}$ (TilS$_{So}$ 8-Ct$_{Sx}$) in *S. oneidensis* WT or *Δhsp90*. The strains WT or *Δhsp90* with plasmid allowing the production of TilS$_{So}$ 8-Ct$_{Sx}$ with hexahistidine tag were grown at 28°C or 34°C and 0.02% arabinose was added. Two hours later, samples were analyzed by Western blot using anti-His antibody. As a control, anti-Hsp90 antibody was used. As control, WT or *Δhsp90* with plasmid allowing the production of TilS$_{So}$ L340P is shown. In *B* and *C*, Western blots shown are representative of two independent experiments. (D) Bacterial growth on liquid media at 28°C of the strains described in *A*. The cells were grown in microplates in LB medium at 28°C with shaking during 24 hours. Data from six replicates are shown as mean ± SD. (E) Bacterial growth on solid media of the strains described in *A.* Cells were grown at 28°C in liquid media and 10-fold serial diluted to be spotted on LB plates. The plates were incubated for 24 hours at 28°C. (F) Bacterial growth on solid media of the strains *S. oneidensis* MR-1 wild-type (WT) or deleted of *hsp90* (*Δhsp90*) carrying *tilS* gene WT or with point mutation allowing the production of TilS with substitution of Leucine 340 to Proline. Cells were grown at 28°C in liquid media and 10-fold serial diluted to be spotted on LB plates. The plates were incubated for 24 hours at 35°C. In *E* and *F*, the plates shown are representative of three independent experiments.

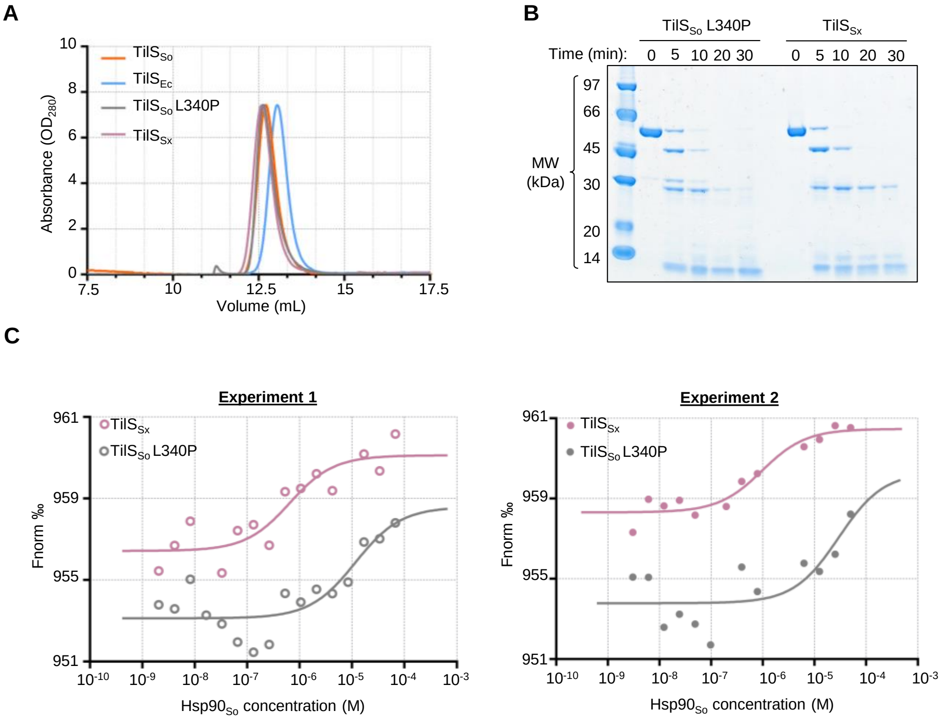


| Protein | $K_d$ (µM) | SD |
|---|---|---|
| TilS$_{So}$ L340P | 20.35 | ± 12.5 |
| TilS$_{Sx}$ | 0.76 | ± 0.23 |

**Figure S9:** ***In vitro*** **characterization of the TilS$_{So}$ single mutant.** (A) Gel filtration analysis of TilS proteins. Purified TilS$_{So}$ WT or L340P, TilS$_{Ec}$, or TilS$_{Sx}$ (8 µM, 100 µL) were analyzed on a Superdex 200 Increase 10/300 GL size exclusion chromatography column. Normalized values of absorbance at 280 nm are shown. (B) Limited proteolysis profiles of purified TilS$_{So}$ L340P or TilS$_{Sx}$ (15 µg) digested with trypsin (0.25 µg) for various times at 37°C in Tris-HCl buffer (50 mM, pH 7.5) containing 100 mM KCl, 10% glycerol and 1 mM DTT in a final reaction volume of 50 µL. At the indicated times, 10 µL was removed and analyzed by SDS-PAGE. (C) Microscale thermophoresis experiment to determine the interaction between Hsp90 and TilS$_{So}$ L340P or TilS$_{EcSx}$. 85 nM of TilS labeled with RED-tris-NTA 2nd generation dye (NanoTemper) was added to serial 2-times dilutions of Hsp90$_{So}$ up to 50 µM. Fluorescence at 670 nm was measured, and values at 2.5 s were used to determine $K_d$. $K_d$ is represented as mean ± SD based on two independent experiments shown on the two panels.

**A**

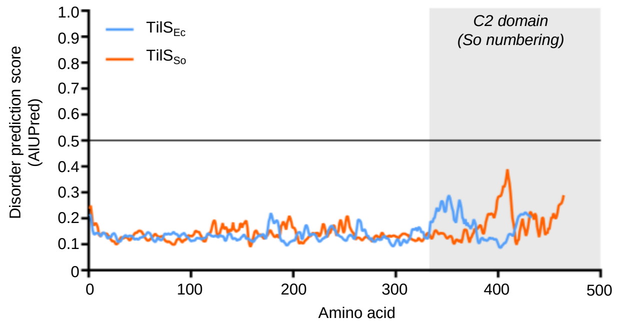


**B**

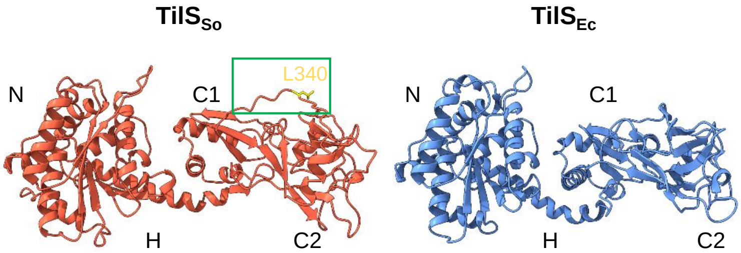


**Figure S10: *In silico* characterization of the TilS$_{So}$ single mutant.** (A) Disorder analysis of TilS$_{So}$ and TilS$_{Ec}$ using AIUPred algorithm (https://iupred.elte.hu/) with the options "AIUPred – Only disorder" and "Default smoothing". The graph shows disorder prediction score by aminoacid. A residue or region showing a score superior to 0.5 can be considered as disordered. (B) Comparison of the structures of TilS$_{So}$ (AlphaFold3 prediction, pTM = 0.81) and TilS$_{Ec}$ (PDB: 1NI5). The green box delineates the region in the vicinity of the leucine residue in position 340 of TilS$_{So}$ (shown in yellow).
